# A comprehensive alignment-filtering methodology improves phylogeny particularly by filtering overly divergent segments

**DOI:** 10.1101/2023.12.26.573321

**Authors:** Qiang Zhang, Xinmei Qin, Yongbin Lu, Pengwei Li, Xiyang Huang

## Abstract

1. Alignment problems may be complicated and diverse and the performance of the existing alignment-filtering tools, particularly their potential effect on phylogenetic reconstruction, have remained debated or limitedly explored.
2. In the present study, we developed a new R package named alignmentFilter to treat the diverse alignment problems, especially masking ambiguously-aligned or overly divergent segments using a newly devised grouping-regrouping algorithms acting on sequence divergence in each of sliding windows throughout alignment. Then we tested and compared the power and accuracy of the prevalent alignment-filtering tools, particularly the effect on phylogeny based on Angiospermae genome-scale data and simulated data.
3. The results indicate the alignment-filtering methods alone may affect the phylogeny decisively, producing (strongly-supported) phylogenetic conflicts. In most cases, alignmentFilter most efficiently improves phylogeny, yielding more congruent phylogeny regardless of phylogeny-reconstruction methods, producing phylogeny more concordant to the priori tree regarding to both topology and branch length in simulation, and most efficiently decreasing the root-to-tip length heterogeneity that is implied to be negatively correlated with topological accuracy.
4. The concepts and algorithms of alignmentFilter are quite distinct from existing alignment-filtering methods. It is not susceptible to specific types of alignment errors that usually baffle other methods and can reach the optimal solution frequently with lowest computation complexity. The biological or statistical sense of the key optional stringency-controlling parameter is more straightforward and the setting and outcome may be more adjustable or predictable. A better and more comprehensive solution of alignment problems in this study will benefit tremendous downstream analyses sourced from alignment and phylogeny.

## 1 INTRODUCTION

Phylogenetic conflict, between/among different data sets as well as various phylogenetic reconstruction methods, is widespread in the tree of life and has been dramatically recognized in the era of genomics. The conflicts were often attributed to, alone or jointly, natural processes such as reticulated evolution (including hybridization as well as horizontal gene transfer) and incomplete lineage sorting of ancestral polymorphism (ILS), stochastic or systematic errors encompassing primary sequence error and methodological deficiency (Lapoint et al., 2011; Mason-Gamer & Kellogg, 1996; Som, 2014; Wong et al., 2007). Previous studies have also suggested possible negative effect of phylogenetic conflict on downstream analyses (e.g. Zhang et al., 2015). This may be especially true if it was not resulted from natural processes or the real cause is not accurately recognized.

Fewer phylogenetic conflict has been recognized as consequence of poor alignment (but see Morrison & Ellis, 1997), though alignment is the cornerstone of downstream phylogenetic, molecular and genomic comparison studies. However, numerous and complicated factors, such as deep divergence, indels, inversion, non-homologous recombination and low-quality assembly of sequence, as well as large data size, may confound aligning (Di Franco et al., 2019). These confront all the presently available aligners, though some of them may outperform others in some situations (Golubchik et al., 2007). Consequently, alignment problem may be more or less inevitable, particularly in the era of genomics when manual inspection of alignment encompassing hundreds to thousands of genes seems unfeasible.

The existing alignment-filtering methods are grounded on distinct concepts and scenarios and may have their own limitations. The block-filtering method, such as Gblocks (Castresana, 2000) and trimAl (Capella-Gutiérrez et al., 2009), identifies and deletes low-quality alignment columns along with probable phylogeny-informative and well-aligned sites, which likely decreases phylogenetic resolution. The segment-filtering method, only known from PREQUAL (Whelan et al., 2018) and HmmCleaner (Di Franco et al., 2019), deletes or masks non-homologous or falsely-aligned string (segment). But both tools were designed for dealing with amino acid (unaligned protein sequences for PREQUAL) or nucleotide codon sequences and may be unsuitable for data of none-coding DNA sequences and/or not applicable for alignment in which problems may be brought later during aligning. Another package TAPER identifies and removes outlier sites through combination of assessing outliers along columns and rows (so-called 2D outlier detection), but it has limitations such as to recognize very short, long and high frequent errors, as noted by the authors (Zhang et al., 2020). Furthermore, all the methods are mainly or solely for handling problematic segment or column, but cannot deal with other diverse alignment problems such as reverse complementary sequence sometimes included during employing programs or pipelines.

The performance of varied alignment-filtering methods and particularly the effect on phylogenetic inference remain inadequately explored or contentious. Some previous studies showed improvement of phylogeny and detection of natural selection site (Castresana, 2000; Di Franco et al., 2019; Jordan & Goldman, 2011; Talavera & Castresana, 2007), but other study demonstrated negative or detrimental effect to phylogeny (Tan et al., 2015). In addition, the possible effect of alignment problem on branch length inference remains far less concerned, though it is expected to be more likely affected as it cannot be overshadowed by the true phylogenetic signals as that for inference of topology. Generally, the degree and implication of possible impact of alignment filtered using various alignment-filtering methods on phylogeny remain ambiguous. Therefore, all these suggest significance and necessity of developing more sophisticated program to address diverse alignment problems and further exploration of the effect on downstream analyses.

In the present study, we developed a R package named alignmentFilter for comprehensive alignment filtration. The power of this newly developed and other prevalent alignment-filtering tools on phylogenetic inference was examined and compared based on both empirical and simulated data. For the empirical data, we focused on the long-standing controversies over the deep angiosperm phylogeny and chose the plastome-scale and transcriptome or genome-scale data from Li et al. (2019) and Yang et al. (2020), respectively. These two studies as well as others show phylogenetic conflicts, particularly among the five Mesangiospermae clades (Gitzendanner et al., 2018; Leebens-Mack et al., 2019; Moore et al., 2007; Qiu et al., 2010; Ruhfel et al., 2014; Soltis et al., 2011; Wickett et al., 2014; Xue et al., 2021; Zeng et al., 2014). The results indicated that alignment-filtering method alone can largely affect inferred phylogeny, and in most cases after alignment filtration by using alignmentFilter both the topological conflict and root-to-tip length heterogeneity are simultaneously minimized most efficiently and the phylogeny in simulation is more consistent with the priori.

## 2 MATERIALS AND METHODS

### 2.1 Concepts and rationales of alignment Filter

In order to solve the diverse problems in DNA alignment, a set of functions were developed. At whole sequence level, the function “revComplement” identifies and adjusts the reverse complementary sequence that may be sometimes included in using phylogenetic pipelines; “alignmentLength” separates alignments according to their lengths; and “anyShortseq” removes short sequence in alignment. At block level, “degap” deletes terminal and embedded gappy sites separately with different stringency. More importantly, at segmental level, “maskSegment” masks ambiguously-aligned or overly divergent segment (collectively called “overly divergent segment” hereafter as they are essentially indistinguishable) based on a newly developed grouping-regrouping process acting on pairwise segment similarity/divergence in each of the sliding windows throughout an alignment.

The basic rationale of “maskSegment” is to identify and remove overly divergent segments which have as low similarity as random alignment to the largest group of recognized well-aligned segments after a grouping-regrouping processes. The calculation of probability of random similarity is under the simplest assumption that the DNA sequences are equally probably composed of the four nucleotides, such as without any GC biases, and do not take any substitution model into account. This may be often violated more or less in realistic data but it, as broadly corroborated in this study, is still expected to behave well in most realistic cases and has the advantage of low computation load. Under the simplest assumption, the probability of random alignment similarity can be formulated as 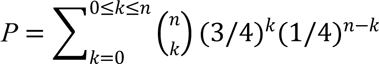, in which, *n* denotes segment length and *k* represents the amount of substitution or variable sites. In a default sliding window with a width of 15bp (*n* = 15), the random alignment similarity (*RS*) can reach at least as high or higher under a given probability (the argument *prob* and its four recommended values in “maskSegment”) is calculated as 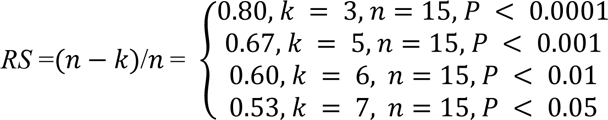. The pairwise similarity (*PS*) are calculated and compared with a corresponding *RS* under a specified probability cutoff value to assess whether they (or the temporal groups they belong to, respectively) should be combined into a same group. The *PS* is similarly but more detailedly formularized as 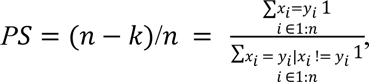, in which x_i_ and y­represent each pair segments in a sliding window, and x_i_ and y_i_ are only compared and counted when they both are definite nucleotides (degenerated and gap sites are skipped). The segments in a sliding window would be initially classified into different temporal groups if the *PS* value lower than the *RS* under the given probability cutoff, meaning statistically significantly falling in the space of random alignment. Subsequently, a regrouping process is then taken between/among the temporal groups after all the sequences had been classified in the first round of grouping. If there is any intermediate sequence in one temporal group that has a *PS* value higher than the *RS* to any sequence in a second temporal group, the two temporal groups are regrouped. This grouping-regrouping process is characterized by not only avoiding unnecessary pairwise comparisons, but also guarantee the best solution. After the grouping-regrouping process, segments other than those in the largest group are finally masked (a flowchart shown in Figure 1). More rationales, tactics and recommended parameter settings are described in Supporting information.

**FIGURE 1.**
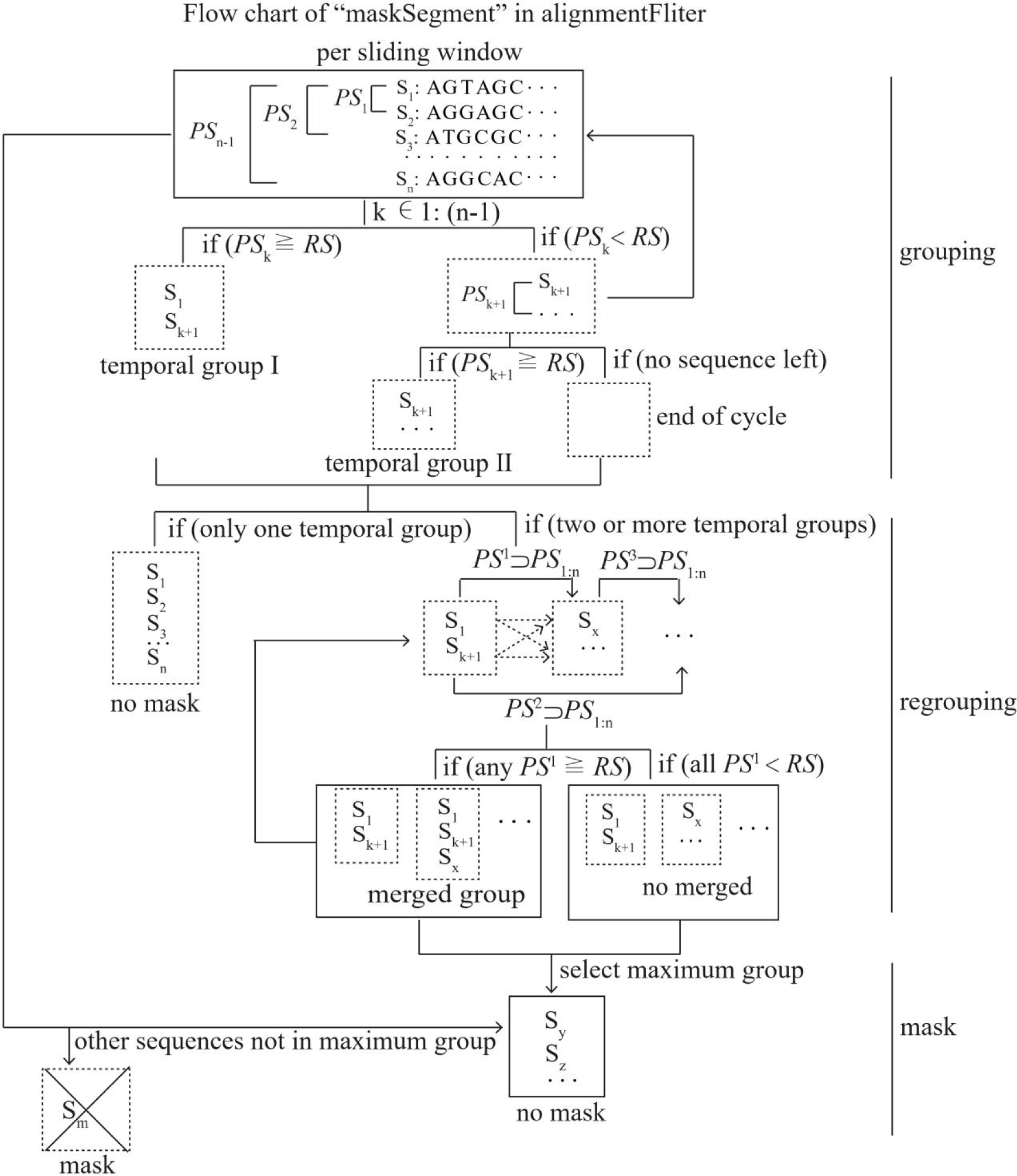
Flow chart of the function “maskSegment”, showing tactics how to isolate and mask divergent segments.

### 2.2 Test effect of alignment filtration on the deep angiosperm phylogenetic conflicts from different data and methods

We downloaded and reanalyzed the plastid data of 80 genes (from 2351 angiosperm species, 353 families and 64 orders) and the data of 296 orthologous nuclear genes (from 151 angiosperm species, 106 families and 51 orders), probably representing densest sampling of angiosperm orders and families until our analyses. The downloaded plastid data had been aligned in the previous study of Li et al. (2019) using MAFFT (Katoh, 2005) and MUSCLE (Edgar, 2004), and the nuclear data had been aligned in the previous study of Yang et al. (2020) using MAFFT. We compared the topological conflicts between data sets as well as between different phylogenetic reconstruction methods, particularly concentrating on whether and how they would be affected by various alignment-filtering methods. The methods including Gblocks, trimAl, HmmCleaner, TAPER and FasParser2 (Sun and Hancock, 2018) were used as comparisons to alignmentFilter.

The parameters were set as follows: probability cutoff (*prob =* 0.05, 0.01, 0.001, 0.00001, respectively, representing increasing stringency and probably masking more overly divergent segments), sliding window size (default *window_width* = 15 nucleotides or 5 codons) and window step increase (default *window_step* = 3 nucleotides or 1 codon) for alignmentFilter; window size for trimming blocks (9 bp), window size for gap regions (9 bp), min out block length (18 bp), number of repeats for sampling (100) for FasParser2; mask symbol as “N” (-m N) for TAPER; type of sequence (-*t* = c) and gap threshold (-b5 = a) for Gblocks; score matrix (costs = −0.15, −0.08, 0.15, 0.45) and symbol fraction (symfrac = 0.50) for HmmCleaner implemented in PhyloSuite (Zhang et al., 2019); similarity threshold (*st* = 0.5, 0.6, 0.7, 0.8, respectively, similarly representing increasing stringency), gap threshold (-*gt* 0.5) and window size (-*w* 7) for trimAl. As very long indels in some plastid alignments may impact phylogenetic reconstruction and not all alignment-filtering tools used contain gap deletion function, the columns with 50 percent or more gaps were uniformly deleted using the “degap” function in alignmentFilter before filtration. All filtering operations ran successfully except FasParser2 on the plastid data, which was too time-consuming to finish.

We reconstructed the phylogeny using RAxML ver. 8.2.12 (Stamatakis, 2014) with GTRGAMMA substitution model and 100 bootstrap pseudo-replicates to calculate node support based on the plastid and nuclear concatenated data, respectively, as well as based on the individual nuclear genes. Concatenation was executed using the function cbind.DNAbin in the R package APE (Paradis et al., 2004). In addition, ASTRAL (Zhang et al., 2018) was further used to infer the species tree based on the nuclear gene trees. The unfiltered data (but the plastid data degapped) was used for phylogenetic reconstruction as comparison too.

To measure the degree of phylogenetic conflict between the plastid and nuclear data and between concatenation-based ML and coalescent-based ASTRAL inferences affected by the different alignment-filtering methods, the (normalized) Robinson-Foulds (RF) tree distances (Robinson & Foulds, 1981) were calculated using the function RF.dist in the R package phangorn (Schliep, 2011). As RF distance calculation requires the same set of taxa for both compared trees, the compared plastid and nuclear trees were reduced using function drop.tip in treeio (Wang et al., 2020) to include the same set of unique orders after renaming the taxa (tips) with the corresponding orders’ names. The reduced concatenation and ASTRAL trees were annotated and visualized using the R package ggtree (Yu et al., 2016). In addition, these trees were further reduced to display the relationships of major Mesangiospermae clades (17 and 16 clades in the plastid and nuclear trees, respectively) which have been argued (Leebens-Mack et al., 2019; Li et al., 2019, 2021; Moore et al., 2007, 2010; Xue et al., 2021; Yang et al., 2020; Zeng et al., 2014, 2017). To more conveniently and intuitively show phylogenetic conflicts caused by the alignment-filtering methods, consensus networks based on each set of the plastid concatenation, nuclear concatenation and nuclear ASTRAL trees were reconstructed using Dendroscope 3 (Huson & Scornavacca, 2012).

### 2.3 Test effect of alignment filtration on root-to-tip length heterogeneity of the nuclear gene trees

The root-to-tip branch length heterogeneity, usually representing molecular evolutionary rate variations among lineages, may be further inflated by false alignment, as aligning problem may likely be clustered in divergent sequences whose root-to-tips are already longer in a tree. Therefore, standard deviation (SD) of the root-to-tip lengths of each tree, was assessed to examine whether and how it would be affected by the various alignment-filtering methods. It can be expected to decrease mostly when possible divergent alignment was effectively adjusted. The sets of (296) individual nuclear gene trees generated from the data sets filtered by the various methods were used, respectively. The trees were rerooted with Amborellales, the earliest diverged angiosperm lineage (Moore et al., 2007; Qiu et al., 1999; Zeng et al., 2014), using function “root” in treeio after all the non-angiosperm taxa were discarded. The root-to-tip lengths per tree was counted using the function node.depth.edgelength in APE and the SD value was calculated using the function sd in R.

### 2.4 Test possible correlation between root-to-tip length heterogeneity of the nuclear gene trees and topological accuracy

Whether the root-to-tip length heterogeneity is correlated with topological accuracy was tested based on the 296 unfiltered nuclear gene trees and the alignmentFilter concatenation ML tree. The gene trees from the unfiltered alignments rather than those based on the filtered data were used because their root-to-tip length heterogeneity may vary more widely enough to show a trend, if existing, and they contain signals of ambiguous alignment whose effect is concerned. The alignmentFilter concatenation ML tree is expected to more likely approach to the true phylogeny than individual gene trees and was also generated using RAxML as the gene trees, so that it was used as the reference tree to which the RF distances of the gene trees were calculated. Similarly, SDs of root-to-tip lengths of the individual gene trees and the normalized RF distances to the reference tree were calculated and their possible correlation was plotted in R.

### 2.5 Test the effect of the various alignment-filtering methods based on simulated data

The above reduced nuclear concatenation ML tree with 53 orders (tips) was used as the priori tree, along which sequences were simulated using the R package PhyloSim (Sipos et al., 2011). A total of 300 genes (alignments) were produced with sequence lengths randomly sampled from 600 to 2,400 bp, tree lengths randomly sampled from 1.02–51.10 (0.1–5 folds of 10.22, the length of the priori tree), and the substitution model GTRGAMMA. For each of the 300 simulated individual genes, five groups with each consisting of two (a total of 10 sequences) out of 53 sequences (tips) were randomly chosen to be partly replaced in each group with a same set of random nucleotide strings, modelling false alignment. For each of the five groups of sequences, five random nucleotide strings with lengths randomly sampled from ranges of 5–10, 11–30, 31–50, 51–100 and 101–400 were produced and used to replace randomly selected segments of the same lengths, rendering the random replacements identical for the two sequences within each group but distinct between groups.

The priori tree, including topology and branch length, was used to benchmark the effect of the various alignment-filtering methods on recovering accurate phylogeny. The same set of methods as used for the empirical data were used to filter the simulated data, respectively. The same phylogenetic inferences were conducted. The RF distances of each set of the concatenation, ASTRAL and individual gene trees to the given priori tree were calculated. The root-to-tip length heterogeneity of individual gene trees was also calculated. Furthermore the degree of reproducibility of true branch length was also assessed by calculating and comparing the mean SD of each root-to-tip length (in proportion to each corresponding tree length) of each of the trees to that of the original priori tree. Although the lengths of the trees used for simulating the 300 genes were scaled with a wide range from 1/10 to 5 folds of the length of the original priori tree, each proportion of root-to-tip to tree length is fixed as the same as that of the original priori tree, against which it is reasonable to compare.

## 3 RESULTS

### 3.1 Different alignment-filtering methods alone produce conflicting deep angiosperm phylogeny with varied RF tree distances

The alignment-filtering methods alone (largely) affect the inferred phylogenetic relationships based on each same set of data and phylogenetic inference method. The sets of inferred plastid concatenation, nuclear concatenation and ASTRAL trees of angiosperm orders and simulation concatenation and ASTRAL trees each show maximum RF distances up to 16, 26, 42, 28, and 12, respectively, suggesting (strong) impact of alignment-filtering methods on phylogenetic relationships. More RF distances between trees from each pair of alignment-filtering methods based on each set of data and phylogenetic methods are listed in Supporting information Tables S1–S5. The conflicting phylogeny (phylogenetic network) is as shown in Figure 2 and Supporting information Figures S1–S4, which are further described in Supporting information.

**FIGURE 2.**
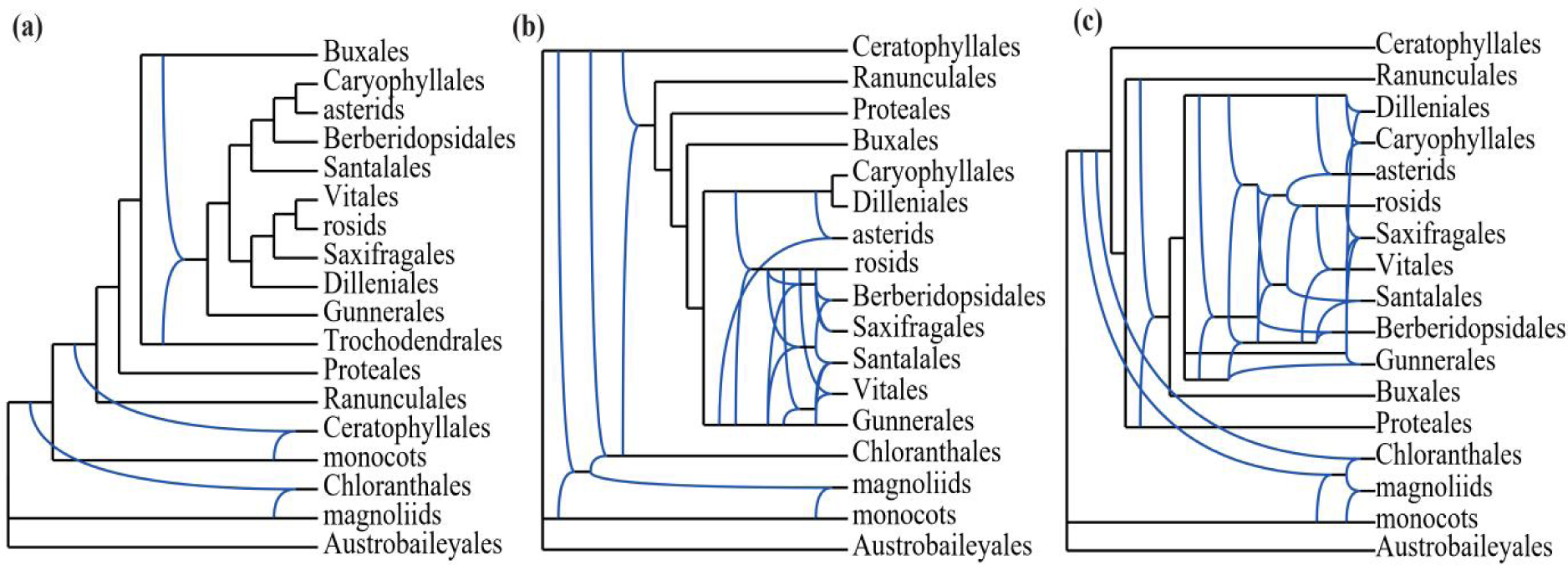
Consensus networks showing phylogenetic conflicts among the major clades in Mesangiospermae invoked by the various alignment-filtering methods. (a) The network of the 11 plastid concatenation ML trees. (b) The network of the 12 nuclear concatenation ML trees. (c) The network of the 12 nuclear ASTRAL trees. The various alignment-filtering methods include alignmentFilter (with parameter of *prob* set to 0.05, 0.01, 0.001 and 0.0001, respectively), FasParser2 (only available for the nuclear data only), Gblocks, HmmCleaner, TAPER and trimAl (with *st* set to 0.05, 0.01, 0.001 and 0.0001, respectively).

### 3.2 AlignmentFilter decreases angiosperm phylogenetic conflict most efficiently

The phylogenetic conflict between the plastid and nuclear data vary among different alignment-filtering methods (Figure 3a). The plastid and nuclear ordinal trees have the lowest RF value 0.38 when using alignmentFilter with *prob* of 0.0001 and trimAl with *st* of 0.7. The second lowest RF distance of 0.41 between the plastid and nuclear concatenation trees is from trimAl with *st* of 0.6. All other methods have the same RF distances of 0.43 between the plastid and nuclear concatenation trees as that based on the unfiltered plastid and nuclear data.

**FIGURE 3.**
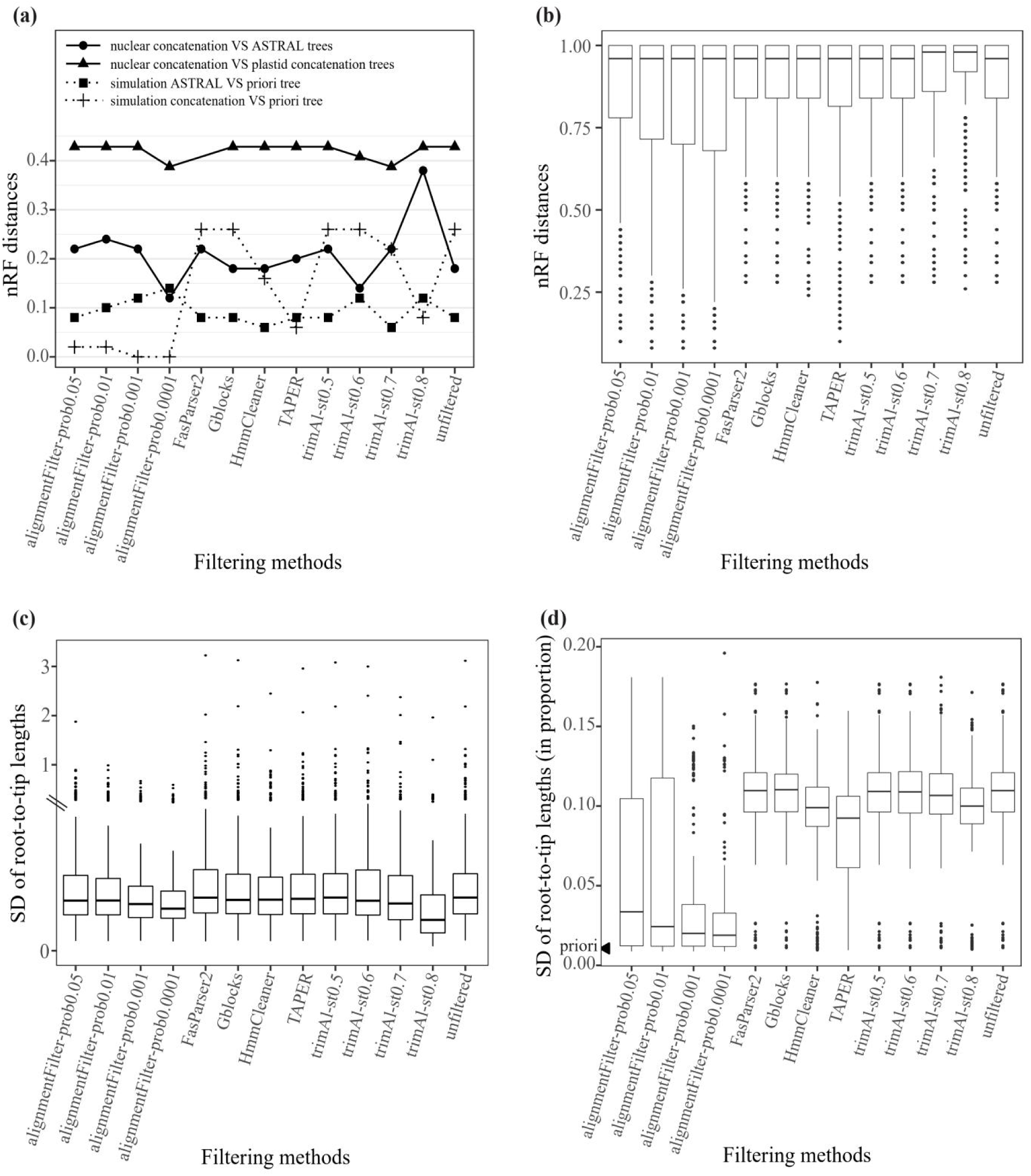
Phylogenetic conflict, topological accuracy and root-to-tip length heterogeneity affected by the various alignment-filtering methods. (a) scatterplot showing that different degrees of phylogenetic conflicts between the plastid and nuclear data, between concatenation-based ML and coalescent-based ASTRAL methods, and different accuracy of the gene trees based on the simulated data to the priori that are all affected by the alignment-filtering methods. (b) Boxplot of the RF distances of each set of the 296 empirical gene trees to the concatenation ML tree across the various alignment-filtering methods, showing alignmentFilter yields gene trees more consistent to the concatenation tree. (c) Boxplot of the root-to-tip length heterogeneity of each set of the 296 empirical gene trees across the various alignment-filtering methods, showing alignmentFilter generally decreases the heterogeneity more efficiently. Note the discontinuity and disproportion on the upper and lower parts of the y-axis for clearer illustration of the boxplot. (d) Boxplot of the root-to-tip length heterogeneity of each set of the 300 simulation gene trees across the various alignment-filtering methods, showing alignmentFilter generally decreases the heterogeneity more efficiently and most close to the priori.

The phylogenetic conflict between the nuclear concatenation and ASTRAL trees have wider variations from 0.12 to 0.38 across the different alignment-filtering methods (Figure 3a). The lowest RF distance 0.12 between the nuclear concatenation and ASTRAL trees is uniquely reached by alignmentFilter with *prob* of 0.0001. The second lowest value 0.14 is from trimAl with *st* of 0.6. All other methods do not minimize the phylogenetic conflict between the nuclear concatenation and ASTRAL trees, with RF values identical to or higher than the value of 0.18 obtained based on the unfiltered nuclear data.

### 3.3 AlignmentFilter trees generally better match the priori tree in topology

In simulation, the concatenation trees are more sensitive to alignment filtration than the coalescent trees, with RF distance ranges of 0–0.26 and 0.06–0.14, respectively (Figure 3a). In concatenation tree inferences, alignmentFilter performs better than all other methods, which (almost) fully recovers the topology of the priori tree, rendering RF distances of 0.02 with *prob* of 0.05 and 0.01 and zero with *prob* of 0.001 and 0.0001, respectively. The suboptimal performance, RF distance of 0.08, is from trimAl with *st* of 0.8. But with less stringent setting, the effect of trimAl dramatically declines, showing RF distances of 0.22 and 0.26, which (almost) reach the value based on the unfiltered data. Another suboptimal performance with RF distance of 0.16 is from HmmCleaner. All the rests have RF distances identical to that based on the unfiltered data.

In the coalescent tree inferences based on the simulated data (Figure 3a), HmmCleaner and trimAl (with *st* of 0.7) perform equally best, both with RF distances of 0.06, slightly lower than that of 0.08 based on the unfiltered data. Gblocks, FasParser2 and alignmentFilter (with *prob* of 0.05) yield the second best RF distance of 0.08 but the same to that based on the unfiltered data. All the rests have RF distances larger than that based on the unfiltered data, with a peak value (0.14) obtained by alignmentFilter with *prob* of 0.0001.

The RF distances of the individual gene trees to the priori tree, particularly the minimum and lower quantile as well as mean RF distances from alignmentFilter decrease (significantly) in contrast to all the other methods (0.08, 0.68 and 0.81, respectively in alignmentFilter, while the respective second lowest values of 0.10, 0.82 and 0.86 achieved in TAPER; Figure 3b). In alignmentFilter, these RF distances generally decrease along with decrease of the parameter *prob*. However, all other methods failed to decrease RF distances of the gene trees to the priori tree, almost having same distribution as that based on the unfiltered simulated data. In trimAl, the RF distances even show a trend of increase along elevation of the parameter *st* and they, with stringent setting, are even worse than that based on the unfiltered data (lower quantile, median and mean reach up to 0.92, 0.98 and 0.92, respectively, for trimAl; while 0.84, 0.96 and 0.89, respectively, based on the unfiltered data).

### 3.4 AlignmentFilter generally decreases root-to-tip length heterogeneity most efficiently

According to the 296 nuclear gene trees, the alignmentFilter decrease the root-to-tip length heterogeneity (denoted by SD values) more efficiently across all methods (Figure 3c). The lowest mean and lowest maximal values (0.13 and 0.59, respectively) are reached by using alignmentFilter with *prob* of 0.0001. Even with lower *prob*, alignmentFilter generally decreases the heterogeneity more efficiently than other methods except timAl with *st* of 0.8, and the heterogeneity, as shown by the four values of mean, median, upper quantile and maximum values, decreases monotonically along with decrease of *prob* in alignmentFilter. By contrast, it is more complex using trimAl with varied *st*, which keep nearly constant except a noticeable decrease with *st* of 0.8. Although trimAl with *st* of 0.8 largely decreases the heterogeneity too (It among all methods yielded the lowest minimum 0.05, lower quantile 0.07, median 0.10 and upper quantile 0.14), it is at the cost of usually degrading topological accuracy (one of examples as shown in Figure 3b). All other methods fail to (significantly) decrease the heterogeneity and show similar distribution of SD values as that from the unfiltered data.

In simulation, the trees from alignmentFilter generally produces lowest root-to-tip length heterogeneity which also most approach to that of the priori tree (Figure 3d). All the best statistic five numbers of the SDs of the root-to-tip lengths of the gene trees are obtained in alignmentFilter, particularly with more stringent setting of *prob* (lowest minimum 0.0087 with *prob* of 0.05, lowest maximum 0.1502 with *prob* of 0.001, lowest lower and upper quantiles, median and mean values of 0.0119, 0.0329, 0.0189 and 0.0287 with *prob* of 0.0001, respectively). TAPER, HmmCleaner and trimAl with *st* of 0.8 decrease the heterogeneity to some extent as well. All the rest methods have heterogeneity distribution of gene trees almost fully overlapped with that based on the unfiltered data.

### 3.5 Correlation between root-to-tip heterogeneity of the nuclear gene trees and their topological concordance to the concatenation tree

The correlation analyses indicated that the RF distances of the nuclear gene trees to the concatenation tree are positively correlated in the main range of the SD values of 0–2 where most individual gene trees locate (Figure 4). It turns to decrease (negative correlation) in the area of SD value larger than 2, where only three genes are situated in, and thus seemingly lack statistic confidence in this outlier region.

**FIGURE 4.**
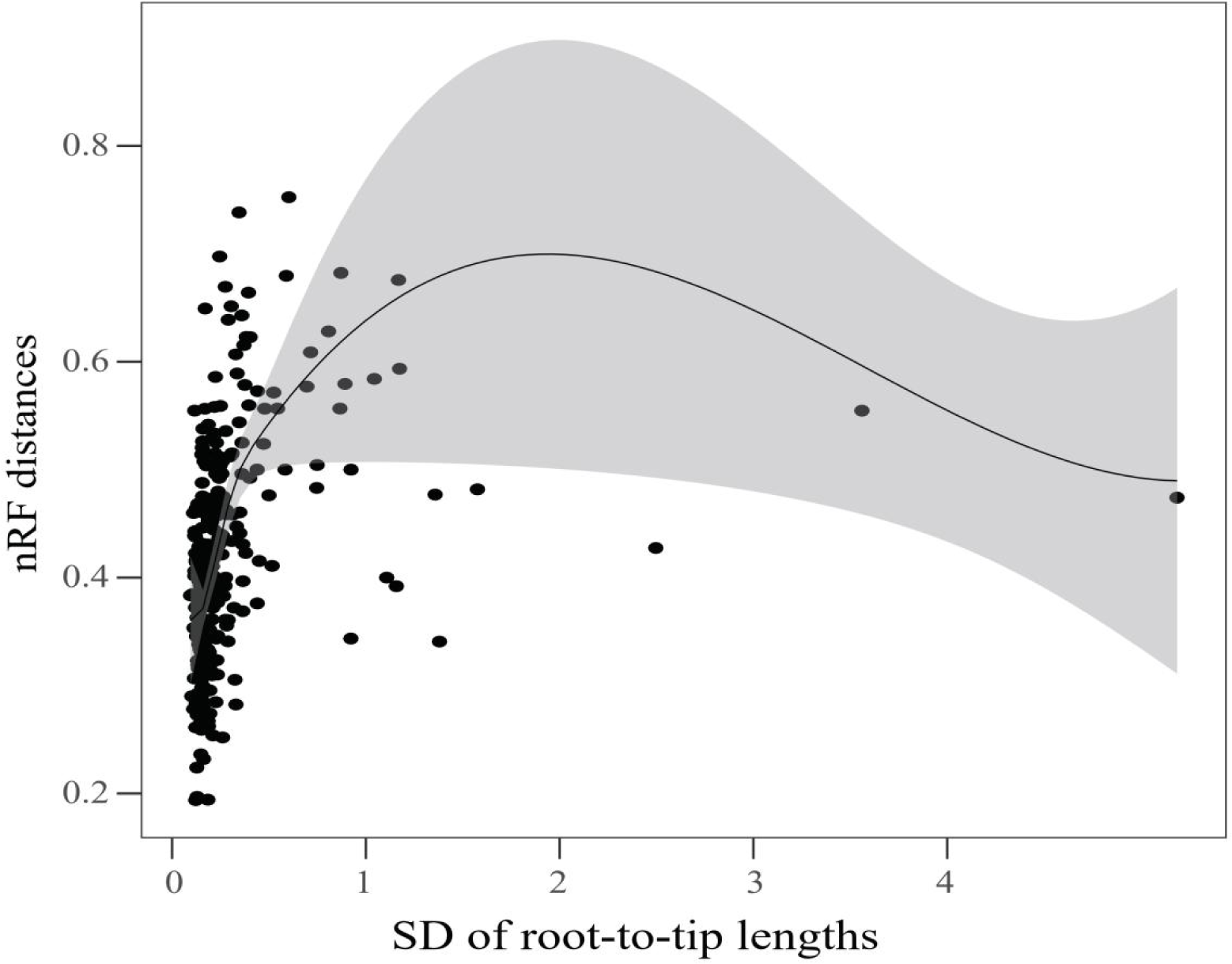
Correlation scatterplot of the root-to-tip length heterogeneity of gene trees and their topological consistency, showing positive correlation at the main region. The main region with SD value lower than ca. 2 contains most samples. The SD of root-to-tip branch lengths of each of the 296 gene trees based on the unfiltered empirical nuclear data and their RF distances to the alignmentFilter (with *prob* set to 0.0001) concatenation ML tree are counted.

### 3.6 AlignmentFilter trees more approximate to the priori tree in root-to-tip length

The trees based on the simulated data filtered using alignmentFilter have root-to-tip lengths significantly more approximating to those of the priori tree (Figure 5). The lowest values of the five numbers of the ranges of root-to-tip length standard deviations of the trees to the priori tree are all from alignmentFilter, particularly with more stringent setting of *prob* (lowest minimum 0.0010 with *prob* of 0.05, lowest lower and upper quantiles of 0.0036 and 0.0285 with *prob* of 0.001, lowest mean, median and maximum values of 0.0188, 0.0133, 0.1007 with *prob* of 0.0001, respectively). In alignmentFilter, the SD values generally decrease monotonically along with decrease of *prob*. The ranges of the SD values of trees from other alignment-filtering methods are strikingly larger than those from alignmentFilter and they, except TAPER, even approach or exceed those based on the unfiltered simulated data.

**FIGURE 5.**
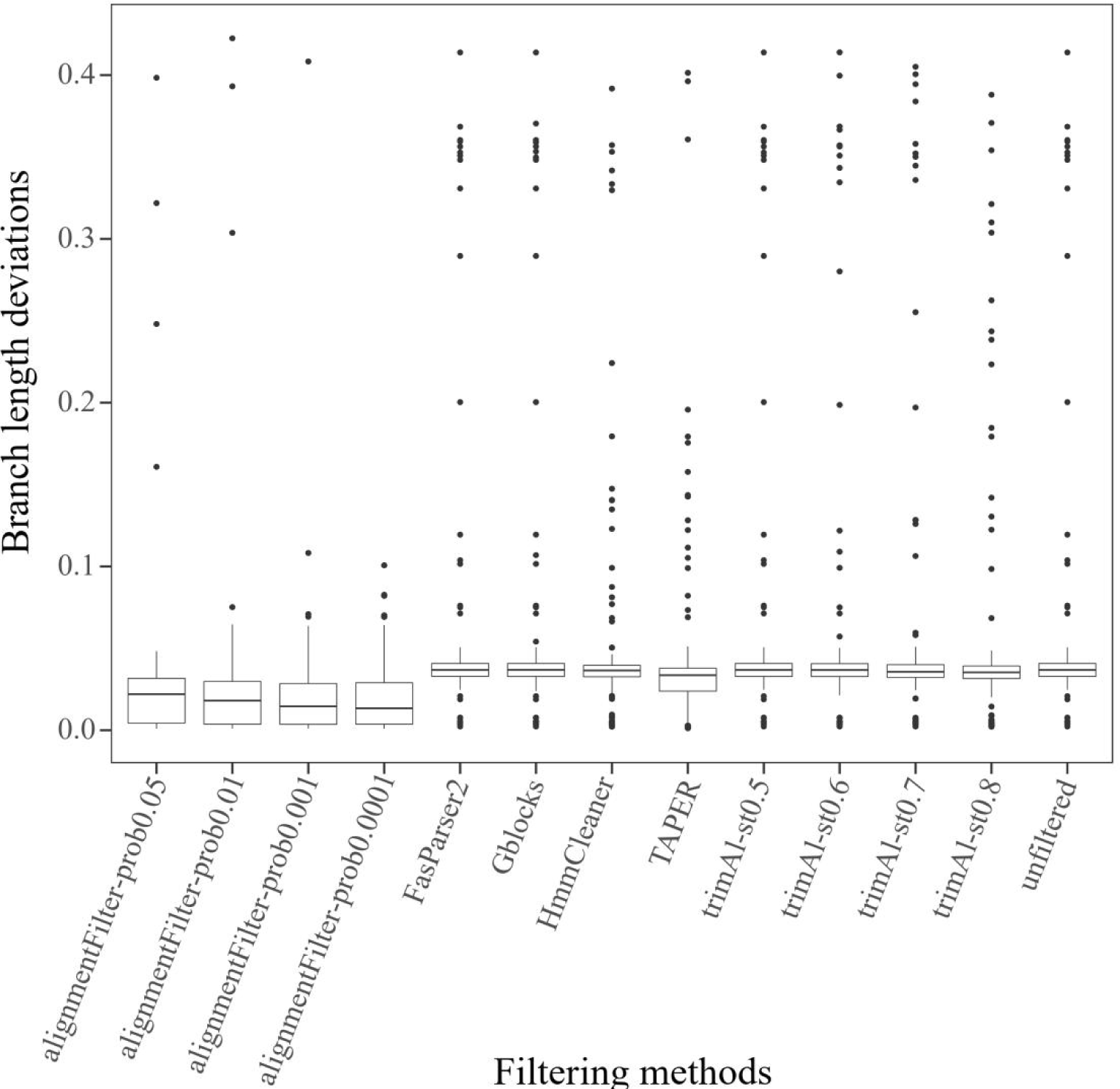
Boxplot of the deviations of root-to-tip lengths from the priori, showing alignmentFilter trees more approximating the priori. Each set of the 300 gene trees based on the simulated data filtered using each of the various alignment-filtering methods are counted.

## 4 DISCUSSION

### 4.1 Overlooked cause of phylogenetic conflict: artifact resulted from divergent alignment along with alignment-filtering methods

Fewer studies have linked phylogenetic conflict with alignment. One of the possible reasons may be due to assumptions that the present aligning and alignment-filtering methods are able to avoid or solve spurious alignment, and/or alignment problem may be less frequent and overshadowed by predominant accurate alignment regions, thus being unable to bias the phylogeny and inducing phylogenetic conflict or artifact. However, our study indicates alignment and alignment-filtering methods indeed can largely affect phylogeny and cause phylogenetic conflict. And even a small proportion of divergent segments masked (ca. 0.05 percent and 1.88 percent sites masked in the plastid and nuclear data, respectively, using alignmentFilter with *prob* of 0.0001) changed the topology and reconciled the phylogenetic conflicts, contradicting the above assumption. These indicate that some phylogenetic conflicts are soft and may be just phylogenetic artifact resulted from poor alignment failed to be appropriately adjusted by the alignment-filtering methods.

If the phylogenetic conflict is artifact, the the exploration of the cause of phylogenetic conflict, including inference of gene flow and ILS, might be biased and false results may be obtained. Therefore, it seems that phylogenetic conflicts widespread in the tree of life and the possible causes might merit re-evaluation, preceded by effort put into effectively solving probable alignment problems whose diversity, solution as well as consequence have been still overlooked, particularly in the era of genomics.

### 4.2 Effect of alignmentFilter on phylogenetic reconstruction and its implications

Phylogeny is in general indicated to be most efficiently optimized by alignmentFilter, particularly with stringent setting of the parameter *prob* and in the nuclear and simulated data sets where there are more alignment problems. The general more consistent deep angiosperm phylogenies from alignmentFilter (Figure 3a) should mean more likely approach to the true evolutionary relationships, as convergence to a same wrong topology is very unlikely due to a gigantic space of possible tree topologies with hundreds to thousands of tips. Even if only five tips are considered, corresponding to the five Mesangiospermae clades, the number of possible rooted tree topologies reaches up to 105. Better reproducibility of true phylogeny, in terms of both topology and branch lengths, was also suggested in simulation analyses, despite an exception of marginal negative effect of alignmentFilter for the ASTRAL inference. This may be occasional and confined to the specific simulated data, such as due to too long simulated random segments in some alignments, which may randomly yield unpredictable topologies due to few phylogenetic signals, despite falsely-aligned segments are accurately masked. Moreover the simulated data with divergent sequences (modelling higher substitution rate) but much less sampling tips in contrast to the empirical data implies lack of intermediate sequences bridging up the large divergence gap and thus is excessively masked, finally deteriorating the phylogeny. This assumption seems to be supported by that the RF distances of the ASTRAL trees to the priori tree increase along with decrease of the parameter *prob* in alignmentFilter as shown in Figure 3a.

Not only the topology but also branch length is improved by alignmentFilter. More accurate root-to-tip length and less heterogeneity across lineages are simultaneously achieved in simulation analyses, and the root-to-tip length heterogeneity of the empirical individual gene trees is efficiently decreased by alignmentFilter as well (Figures 3c, 3d and 5). Although it is unknown whether lower heterogeneity among lineages really approximate the true value for the empirical data, it seems to be beneficial as the heterogeneity is negatively associated with topological accuracy, showing a trend of more congruence to the concatenation tree of the gene trees with lower heterogeneity (Figure 4). This correlation may be due to that heterogeneity and topology conflict are both linked with false alignment. And heterogeneity even if purely resulted from lineage-specific molecular evolutionary rate variations may affect inference of phylogenetic relationship too, such as via long branch attraction from accelerated substitution. The benefit of decreased heterogeneity may be extended to other downstream analyses, such as molecular dating and phylogenetic diversity estimate, since they require properly adjusting molecular evolutionary rate variation too (Zhang et al., 2015).

The performance and effect of alignmentFilter has been extensively tested and is expected to work well in more diverse conditions. The empirical angiosperm-scale plastid, nuclear and simulated data encompass large variations of individual gene length, substitution rate (50 folds between the lowest and highest), sampling density of taxa (thousands, hundreds to dozens of accessions), and ratio of poor alignment (from few in empirical data to ten out of 53 sequences in the simulated data, and from several sites to hundreds of contiguous sites in a sequence). Performance of combination of the various alignment-filtering and phylogenetic reconstruction methods has been widely compared as well. AlignmentFilter also performs well to the data of very short, long and high frequent alignment errors that are usual limitations to TAPER (Zhang et al., 2020) and other methods as well according to our observation (Supporting information Figure S5). Nonetheless, there remain unexplored spaces deserving future investigation. It includes but is not limited to untangle the impact of more diverse data of distinct characteristics, such as that characterized by varied tree topological structure (for example differential topological symmetry) and larger substitution rate heterogeneity, suitability of parameter settings, and performance of combination with different upstream aligning tools, such as MUSCLE (Edgar, 2004) and MACSE (Ranwez et al., 2018), and effect on more other downstream analyses.

### 4.3 The long-debated deep angiosperm phylogeny

The relationships of Ceratophyllales sister to eudicots and Chloranthales sister to magnoliids are recovered in both plastid concatenation and nuclear ASTRAL trees in this study using alignmentFilter (with *prob* set to 0.0001; Figures 6 and 7). These results somewhat reconcile the conflicts obtained in the previous studies based on the two datasets (Li et al., 2019; Yang et al., 2020). The relationships of the five Mesangiospermae clades in the alignmentFilter plastid tree coincide with those from the plastid phylogenomic study of green plants (Gitzendanner et al., 2018). Moreover, the Mesangiospermae relationships in the alignmentFilter nuclear ASTRAL tree (with *prob* of 0.0001) are the same as those in the recent phylogenomic studies of green plants based on 1,000 transcriptomes (Leebens-Mack et al., 2019) and Mesangiospermae based on genomic data but much sparser sampling of taxa (Guo et al., 2021; Ma et al., 2021). The more convergent relationships achieved suggests robustness and both the intergenomic and intragenomic Mesangiospermae phylogenetic conflicts seem not so wild as shown in the previous studies (Leebens-Mack et al., 2019; Li et al., 2019; Moore et al., 2007; Yang et al., 2020; Zeng et al., 2014). However, monocots is usually suggested to be more close to eudicots or eudicots plus Ceratophyllales lineage, with low to moderate support, in this (BS = 87) and previous studies based on plastomes (Gitzendanner et al., 2018; Li et al., 2019; Moore et al., 2007) or more close to magnoliids based on genomic microsynteny (Zhao et al., 2021). In addition, ancient gene flow or ILS suggested to be prevalent in diverse rapidly-diversified organisms (García et al., 2017; Hou et al., 2022; Koblmüller et al., 2010), were also identified among the deep Mesangiospermae clades (Guo et al., 2021; Ma et al., 2021). Therefore, the cause of the minimized but persisting conflicts still deserves further exploration.

**FIGURE 6.**
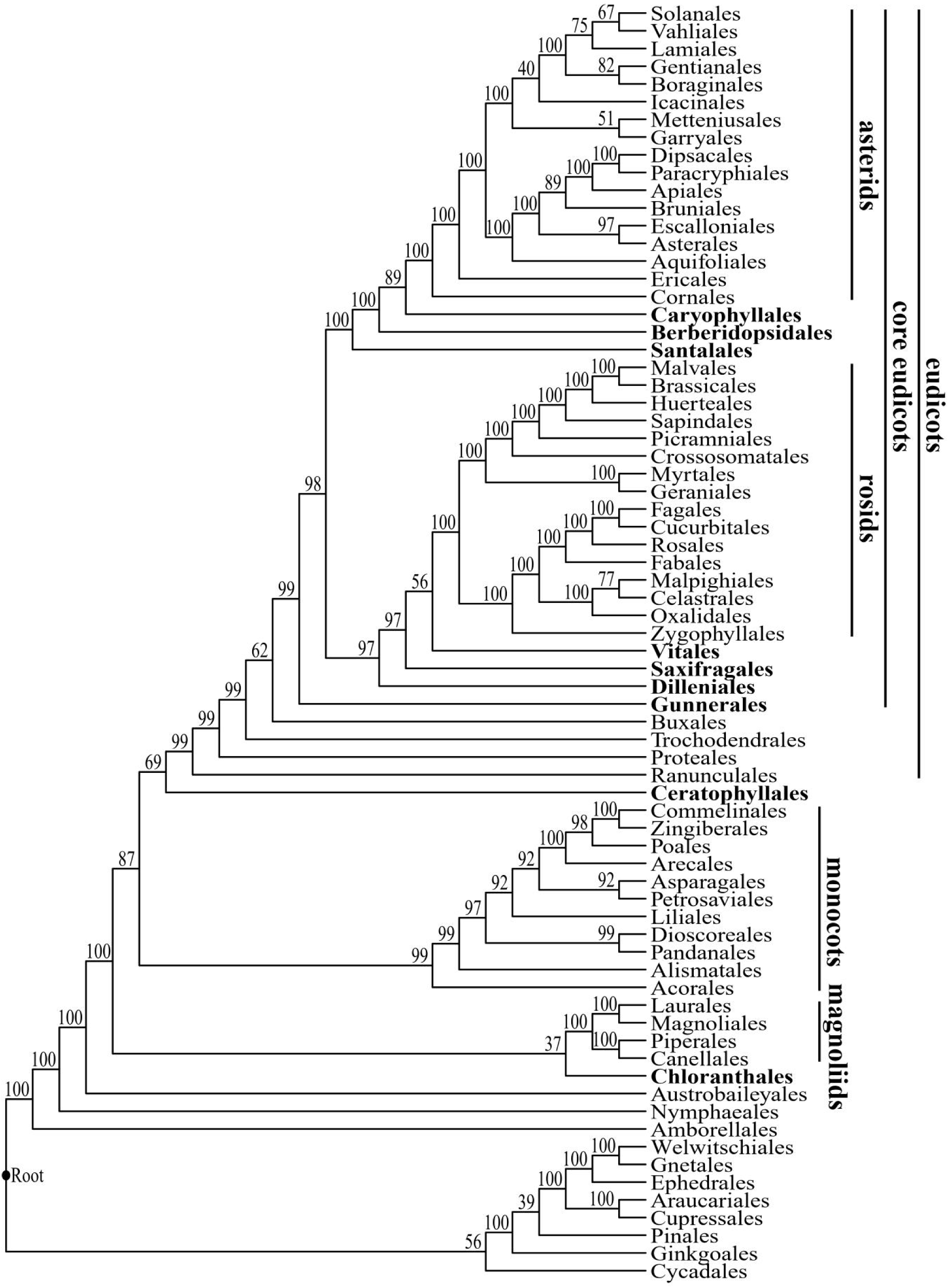
The reduced plastid concatenation ML tree from alignmentFilter (with *prob* of 0.0001), showing the phylogenetic relationships of angiosperm orders. Branch support values were obtained from 100 bootstrap analyses. The key higher taxonomic units, i.e. magnoliids, monocots, eudicots, core eudicots, rosids and asterids are annotated and other orders with concerned phylogenetic positions are highlighted in bold.

**FIGURE 7.**
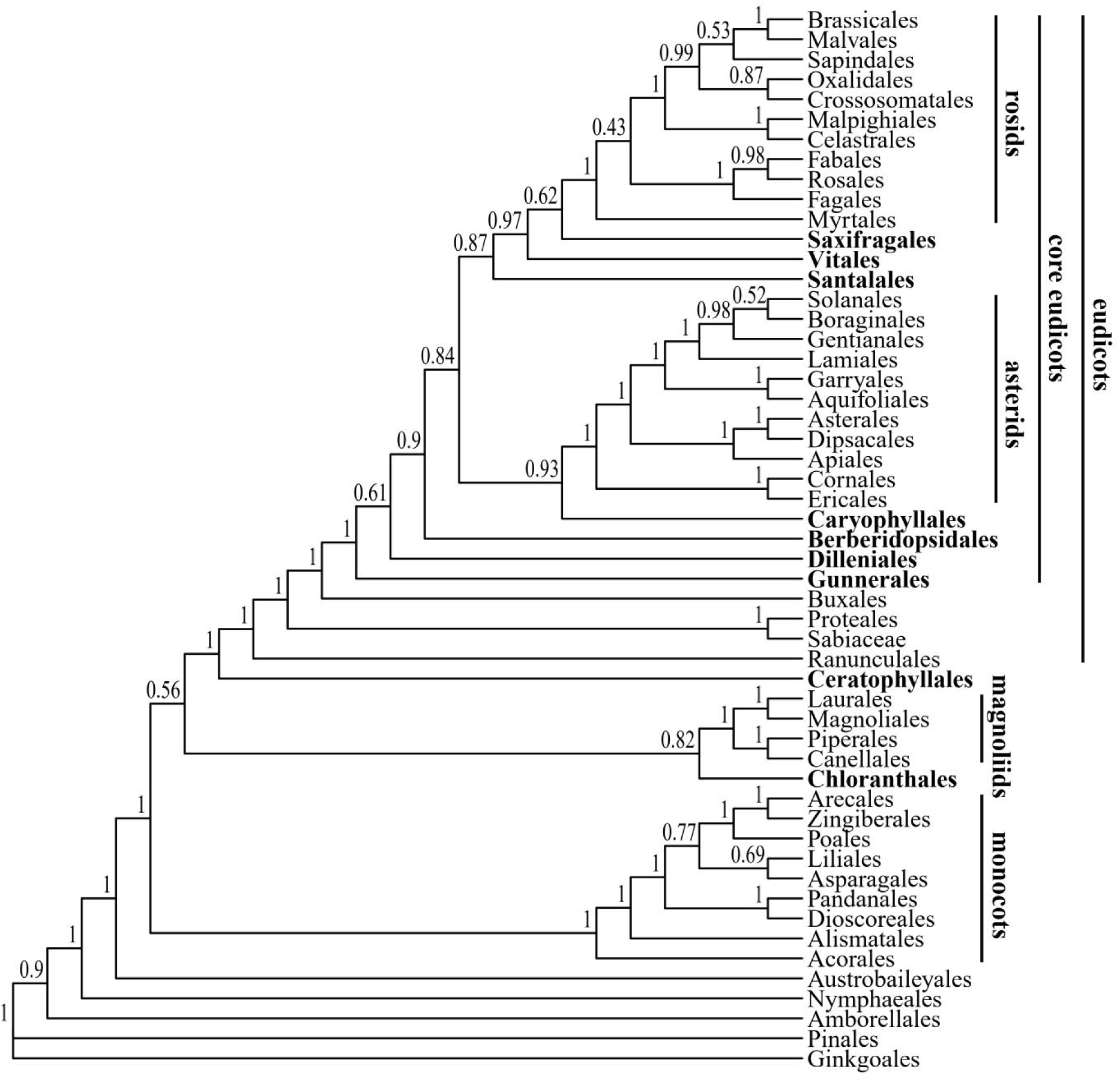
The reduced nuclear ASTRAL tree from alignmentFilter (with *prob* of 0.0001), showing the phylogenetic relationships of angiosperm orders. Local posterior probability values are given to each branches. Key higher taxonomic units and orders with concerned phylogenetic positions are annotated or highlighted as in Figure 6.

More phylogenetic conflicts exist among the core eudicot clades. Only the clade consisting of rosids, Saxifragales and Vitales and the sister relationship of asterids and Caryophyllales are predominantly recovered in both the plastid concatenation and nuclear (ASTRAL) trees in this as well as the previous studies (Li et al., 2019; Zeng et al., 2017). The phylogenetic placement of the other core eudicots is still variable between the plastid and nuclear trees and even between the nuclear concatenation and ASTRAL trees (Figures 6–8). ILS may be a possible cause of the conflicts, as the deep core eudicots rapidly diversified in Early Cretaceous (ca. 125 Ma; Zeng et al., 2017) and conflicts exist between nuclear concatenation and ASTRAL trees. Wrong or conflicting phylogeny may be inferred when there are high level of ILS, falling in the so-called anomaly zone (Mendes & Hahn, 2018). In addition, the ancestor of core eudicots as a whole experienced an ancient genome triplication coupled with rapid radiation (Jiao et al., 2012), which may confound accurate identification of orthology of nuclear genes and distort the phylogeny (Xiong et al., 2022). These hypotheses and whether other processes such as hybridization or gene introgression are also involved in the conflict need validation and in-depth exploration.

**FIGURE 8.**
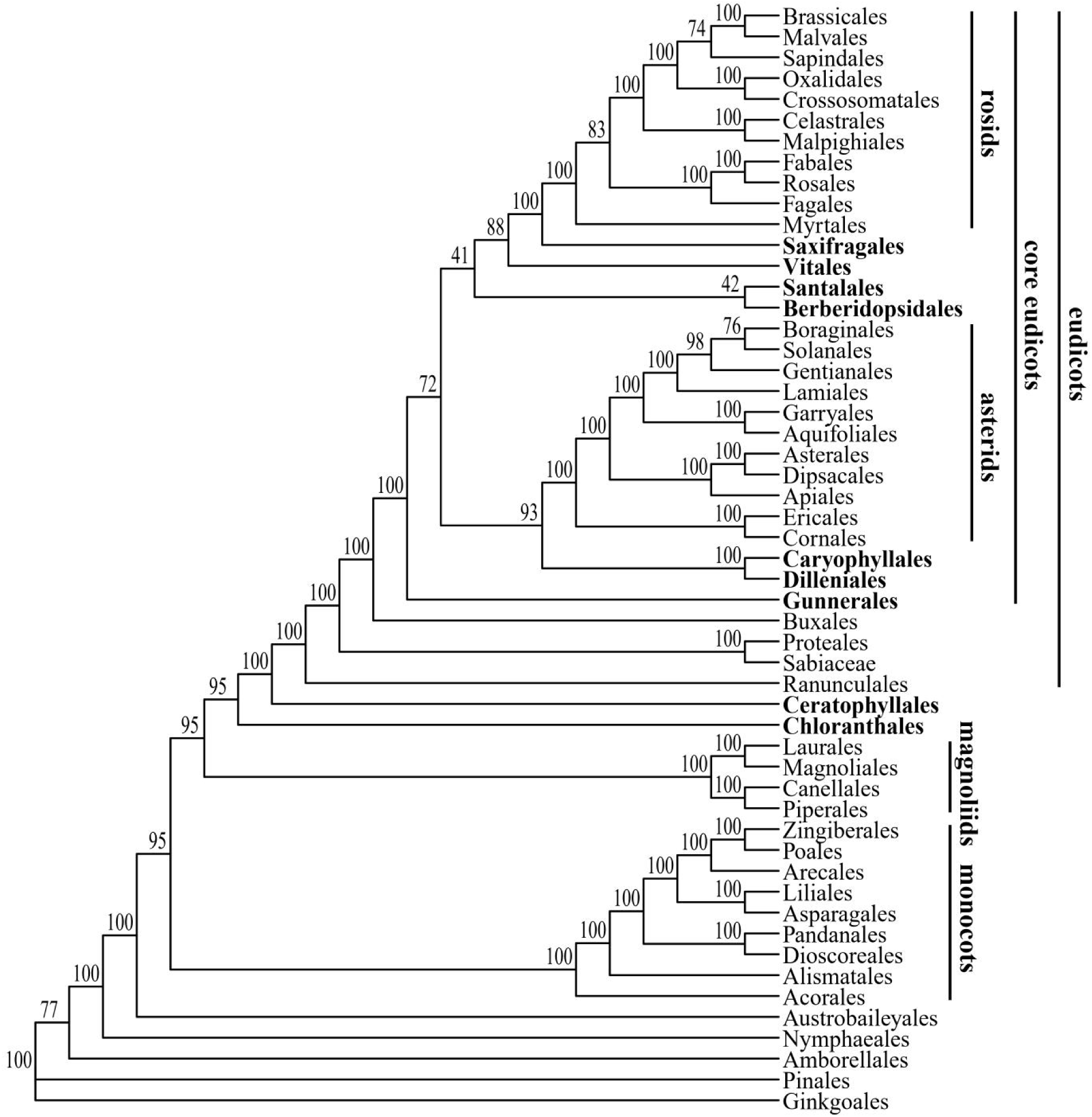
The reduced nuclear concatenation ML tree from alignmentFilter (with *prob* of 0.0001), showing the phylogenetic relationships of angiosperm orders. Branch support values were obtained from 100 bootstrap analyses. Key higher taxonomic units and orders with concerned phylogenetic positions are annotated or highlighted as in Figure 6.

## SUPPORTING INFORMATION

Supporting information includes experimental procedures, results, references and items (Tables S1–S5; Figures S1–S5).

## AUTHOR CONTRIBUTIONS

Qiang Zhang and Xiyang Huang designed the study and drafted the manuscript; Qiang Zhang wrote the program; Qiang Zhang and Xinmei Qin performed data analyses, Qiang Zhang, Xinmei Qin, Yongbin Lu and Pengwei Li revised and improved the manuscript.

## ACKNOWLEDGEMENTS

This work was supported by National Natural Science Foundation of China (grant number 32260045), Guangxi Natural Science Foundation (grant number 2021GXNSFAA196049), National Natural Science Foundation of China (grant number 31660084), and Basic research fund of Guangxi Academy of Sciences (grant number CQZ-C-1901). We thank Bin Wang from Guangxi Institute of Botany for help solve a technique problem when building the package.

## DATA AVAILABILITY STATEMENT

AlignmentFilter is freely available from https://github.com/qiangzhang04/alignmentFilter, and the code and other data produced in this study will be uploaded to Dryad Digital Repository soon before publication. The empirical plastid and nuclear data from Li et al. (2019) and Yang et al. (2020) is downloaded from the Dryad Digital Repository (,https://doi.org/10.5061/dryad.bq091cg) and the figshare (,https://figshare.com/s/27c41bba65a30dbfd3c7), respectively.

