## supplementary text,tables and figures for "A comprehensive alignment-filtering methodology improves phylogeny particularly by filtering overly divergent segments"

**Supporting information**

**Experimental procedures**

**Concepts and rationales of alignmentFilter.**—The parameters controlling the sliding window, i.e. window and step increase sizes (*window_width* and *window_step*, respectively), are optional with default of five codons (or 15 sites) and one codon (three sites) for coding gene sequences (or other none-coding nucleotide sequences), respectively. The *PS* value is formularized as $PS = \frac{\sum_{x_i=y_i,\ i\in 1:n} 1}{\sum_{x_i=y_i | x_i != y_i,\ i\in 1:n} 1}$, in which $x_i$ and $y_i$ represent each of paired segments in a sliding window with length *n*, and $x_i$ and $y_i$ are only compared and counted when they both belong to definite nucleotides (degenerated and gap sites are treated as missing). So that, the *PS* represents the proportion of the number of pairwise matched residues to the total number of all compared pairwise residues, with gap and degenerate sites, if there exist, not taken into account. The segments in a window will be classified into (different) temporal group(s) according to the similarity scores of each of pairwise segments when *PS* is as low or lower than random similarity (*RS*) under a give probability cutoff (the parameter *prob*), cycling the process with $x_i$ assigned with the first segment (remained) and $y_i$ sequentially represented by each of all the other segments remained. With the cycle going on, the segments are continually classified into temporal group(s). After all the segments in a sliding window are classified into temporal group(s), a new round of grouping process is initiated to further merge any two temporal groups if there is any segment from one group that has *PS* value over the *RS* with any segment from other groups (a grouping-regrouping tactic). It is notified that if a pair of segments have less than three paired definite nucleotides compared, they would be classified into isolated group(s) to avoid false grouping by high likely random similarity. After all these processes end, it would turn out that within each group, the segments have *PS* over *RS* with each other, or they respectively have at least one *PS* values over *RS* to a same third segment; while segments from different groups have all pairwise *PS* scores as low or lower than *RS* and none segment from one group that is available to interconnect any other group with a *PS* higher than *RS* with any segment from other groups. The sliding window moves across the entire columns of alignment. If the segments must be classified into at least two groups, the smaller group(s) consisting of less number of segments is masked with degenerate base "N" for the sites while the (one of) largest group remains intact in the window.

The grouping-regrouping tactic newly devised in alignmentFilter is strikingly distinct from all other alignment-filtering methods, which has an advantage to obtain the optional solution most often without significantly increasing computation load. The computation time is linear with amount of segment pairwise comparisons. Existing analogous methods usually achieve optional solution by exhaustive comparisons between all pairwise segments, which have a computation burden of $n(n\text{-}1)/2$ in a sliding window with $n$ denoting total number of sequences. This heavy burden can decrease as lowest as to $n$-1 comparisons if the first segment has similarities all above the cutoff to the rests and it would not largely increase except predominant sequences are overly divergent between each other. In a correct alignment that includes divergent but true homologous sequences, such as those spanning higher taxonomic ranks or experiencing accelerated substitution rates may be masked by existing methods. AlignmentFilter otherwise may somewhat rescue this via grouping-regrouping processes if it is of dense sampling, such that intermediate sequences (segments) are likely included which can bridge up the divergence gaps, linking them into a same group and avoiding excessive filtration. Additionally, the rationale/criterion purely depending on sequence divergence/similarity is parsimonious and easily understandable and the outcome may be more predictable with different levels of stringency of setting the parameter *prob*.

Some other related functions controlling alignment quality are also provided. Reverse complementary sequence may be sometimes included in using the commonly-used pipelines, so that the function “revComplement” was developed to identify and make the corresponding reverse complementary sequence automatically based on input files of sequence data and blast result, the latter of which is produced by using the tool BLAST+ downloaded from NCBI with one of the sequences as query and the whole sequences as the database. The function “alignmentLength” is used for separating the matrices according to the lengths, as short matrices may be less reliable such as for deciding orthology and thus affect phylogenetic inference. The function “anyShortseq” is used for identifying individual short sequence(s) contained in alignment, as significant shorter sequence(s) included in alignment may affect orthology decision, aligning and phylogenetic inference. The terminal gaps (beginning from the first site or ending at the last site of an alignment) are permitted to be assessed and removed at first by setting the parameter *p_flankgap* and then all the rest gappy sites are assessed by setting another parameter *p* in the function “degap”. The independent settings of the two parameters are according to our experience that the terminal gaps may more likely contain sequencing and/or aligning errors that need more stringent cutoff value than for the embedded gaps.

**Setting of the optional parameters in alignmentFilter.**—The key parameter *prob* controlling the stringency is optional and able to be freely specified by the users according to their prior knowledge about the data. In common sense, sequences across higher taxonomic taxa such as families, orders, classes may indeed contain sequences of larger divergence than that from lower taxonomic units such as genus. And non-coding sequences usually evolve more rapidly and are of larger divergence than genes. Higher *prob* (less stringency, e.g. 0.05) may be expected to be more suitable for these true divergent sequences. Otherwise segments of true homologue may be excessively masked particularly when true divergence is in addition to sparse sampling, and thus the remaining very conservative regions are unable to provide enough phylogenetic information. However setting of lower *prob* may conventionally perform better, even if on angiosperm-scale data with dense sampling as suggested in this study. Therefore, more stringency with lower *prob* setting is recommended, particularly for concatenation-based phylogeny inference, until excessive masking begins to degrade phylogenetic reconstruction.

For setting of other parameters, increase of the width (*window_width*) and step size (*window_step*) of a sliding window would decrease calculation burden and speed up but could be less effective for short ambiguous segment in alignment. By contrast, decrease of these parameters may be beneficial for identifying very short ambiguous segment, but risk to mask true phylogenetic informative sites. In fact, to find very short ambiguous segment is expected to be fulfilled by decreasing the parameter *prob* with default settings of others. Therefore, it is recommended to use default settings for these parameters, which is expected to perform commonly better as indicated in this study, except necessities for very special cases.

### Results

**Different alignment-filtering methods alone produce variable conflicting deep angiosperm phylogeny.**—The ML trees and deduced consensus network based on the plastid data filtered by the various alignment-filtering methods show somewhat conflicting phylogenetic relationships among the five Mesangiospermae clades (Figure 2a; Supporting information Figures S1a0 and S2). The tree based on the degapped but unfiltered plastid data shows Ceratophyllales is sister to eudicots (BS = 69), successively followed by monocots (BS = 82) and a clade (BS = 37) consisting of magnoliids and Chloranthales. These relationships were also recovered by using alignmentFilter (BS = 64, 87, 40; 73, 77, 36; 79, 88, 37; 69, 87, 37 for these three nodes, respectively, with the parameter *prob* from 0.05 to 0.0001) and TAPER (BS = 84, 99, 43). The Gblocks, HmmCleaner and trimAl (with the parameter *st* to 0.5) trees all show Chloranthales is grouped with the concordant clade (BS = 78; 83; 68) consisting of eudicots and Ceratophyllales (BS = 60; 98; 89), followed by monocots (BS = 66; 79; 64), and magnoliids split first. By contrast, trimAl with *st* to 0.8 generated relationships that eudicots is sister (BS = 61) to a clade consisting of monocots and Ceratophyllales (BS = 83), which in turn are successively grouped with Chloranthales (BS = 73) and magnoliids. All these results obtained in the present study contradicts more or less with that of Li et al. (2019), in which Chloranthales, magnoliids, monocots, Ceratophyllales and eudicots successively split from the remaining taxa (BS = 29, 72 and 65, respectively).

The nuclear concatenation ML trees and deduced consensus network from different alignment-filtering methods show distinct phylogenetic relationships among the five Mesangiospermae clades (Figure 2b; Supporting information Figures S1b0 and S3). The ML tree based on the unfiltered nuclear data shows Chloranthales is grouped with eudicots (BS = 100) and monocots is sister to magnoliids (BS = 100). The two clades are grouped together (BS = 53) with Ceratophyllales split first. This topology was recovered by using Gblocks (BS = 100, 100, 45), HmmCleaner (BS = 100, 100, 55), TAPER (BS = 100, 100, 61) and alignmentFilter with *prob* set to 0.05 (BS = 100, 100, 48). The trees of FasParser2 (Sun and Hancock, 2018), alignmentFilter with *prob* set to 0.01 and 0.001 and the trimAl with *st* set to 0.5 show that Chloranthales is grouped with eudicots first (BS = 100; 100; 90; 100), which is successively followed by magnoliids (BS = 64; 61; 64; 45), monocots (BS = 100; 100; 100; 100) and finally Ceratophyllales. The trees of alignmentFilter with *prob* set to 0.0001 and the trimAl with *st* to 0.6, 0.7 and 0.8 show Ceratophyllales is sister to eudicots (BS = 100; 47; 99; 64), which is successively followed by Chloranthales (BS = 95; 47; 99; 60), magnoliids (BS = 95; 47; 99; 62) and finally monocots.

The nuclear ASTRAL trees and deduced consensus network also show variable phylogenetic relationships among the five Mesangiospermae clades (Figure 2c; Supporting information Figures S1c0 and S4). The ASTRAL tree based on the unfiltered nuclear data shows magnoliids is sister to monocots (local posterior probability, LPP = 1.00) and they diverged from the clade (LPP = 0.98) made up of the rest three, in which, Ceratophyllales is sister to eudicots (LPP = 0.49) with Chloranthales split first. This topology is the same as that in most other nuclear ASTRAL trees including of alignmentFilter with *prob* set to 0.05 (LPP = 1.00, 0.99, 0.38), 0.6 (LPP = 1.00, 0.98, 0.49) and 0.7 (LPP = 1.00, 0.88, 0.87), FasParser2

(LPP = 1.00, 0.99, 0.39), Gblocks (LPP = 1.00, 0.99, 0.58), HmmCleaner (LPP = 1.00, 0.98, 0.63), TAPER (LPP = 1.00, 0.98, 0.58), trimAl with *st* to 0.5 (LPP = 1.00, 0.98, 0.53) and 0.6 (LPP = 1.00, 0.98, 0.47). But the trimAl ASTRAL tree with *st* to 0.8 shows magnoliids is sister to Chloranthales (LPP = 0.98), they are further grouped with monocots (LPP = 1.00), and Ceratophyllales is sister to eudicots (LPP = 0.93). However, the ASTRAL trees of alignmentFilter with *prob* set to 0.0001 and the trimAl with *st* set to 0.7 show a same topology for the five Mesangiospermae clades, in which, Ceratophyllales is sister to eudicots (LPP = 1.00; 1.00), Chloranthales is sister to magnoliids (LPP = 0.82; 0.89), and the two clades are grouped together (LPP = 0.56; 0.55) with monocots split first. They are different from all above ASTRAL trees but are consistent with a few recent phylogenomic studies based on nuclear data with distinct sampling of taxa and genes (Guo et al., 2021; Leebens-Mack et al., 2019; Ma et al., 2021).

For the phylogenetic relationships among the major clades in eudicots and core eudicots, all the plastid ML trees regardless of the alignment-filtering methods used show congruent phylogenetic relationships, except among Buxales, Trochodendrales and core eudicots (Figure 2a; Supporting information Figures S1a0 and S2). Trochodendrales is suggested to be sister to core eudicots (BS = 40; 45; 62; 43) by using Gblocks and trimAl with *st* to 0.6, 0.7 and 0.8, while all the rest results show Buxales is sister to core eudicots with weak to moderate support (BS = 57; 66; 67; 62; 71; 64; 49 for alignmentFilter with *prob* set to 0.05–0.0001, HmmCleaner, TAPER and trimAl with *st* set to 0.5, respectively).

All the nuclear concatenation ML trees and deduced network except two trimAl trees show consistent relationships among the major eudicots and core eudicot clades (Figure 2b;

Supporting information Figures S1b0 and S3). In core eudicots, Gunnerales split first from the rests which in turn diverged into two clades. In one clade, Berberidopsidales plus Santalales, Vitales, Saxifragales and rosids sequentially diverged from the rest(s). In the other clade, Caryophyllales is sister to Dilleniales and they are grouped with asterids. Most of the branches are moderately (BS = 80–90) to strongly (BS > 90) supported. But the trimAl tree with *st* to 0.7 shows different phylogenetic relationships among Berberidopsidales, Santalales and Vitales, in which Santalales is sister to Vitales (BS = 78) rather than sister to Berberidopsidales. Another trimAl tree with *st* to 0.8 shows more distinct phylogenetic relationships. A clade consisting of Dilleniales and Caryophyllales (BS = 100) diverged from the rest core eudicots. The rest core eudicots (BS = 99) split into two clades. In one clade, rosids is sister to a weakly supported subclade (BS = 21). The subclade further split into two weakly supported groups with one consisting of Saxifragales and Berberidopsidales (BS = 87) and the other (BS = 12) comprising Vitales and a pair of unexpected sisters (BS = 90) made up of Santalales and Gunnerales. The other clade includes asterids only.

The nuclear ASTRAL trees from different alignment-filtering methods also show congruent relationships for basal eudicots (with a rare exception that Proteales rather than Ranunculales diverging first from the rest eudicots in the trimAl tree with *st* set to 0.8), but more variable phylogenetic relationships among the major core eudicot clades (Figure 2c; Supporting information Figures S1c0 and S4). The ASTRAL tree based on the unfiltered nuclear data shows Gunnerales (LPP = 0.53), Dilleniales (LPP = 0.90) and Berberidopsidales (LPP = 0.98) successively diverged from the rest core eudicots. The rest core eudicots split into two clades. One clade includes Caryophyllales and asterids (LPP = 0.74). In the other clade (LPP = 0.87), Santalales (LPP = 0.97), Vitales (LPP = 0.89), Saxifragales (LPP = 1.00) and rosids sequentially diverged from the rest(s). This topology is the same as that in the ASTRAL trees of alignmentFilter with *prob* set to 0.0001 (LPP = 0.61, 0.90, 0.84, 0.93, 0.87, 0.97, 0.62, 1.00) and Gblocks (LPP = 0.71, 0.82, 0.98, 0.73, 0.71, 0.97, 0.77, 1.00). The ASTRAL trees of HmmCleaner, trimAl with *st* set to 0.5 and alignmentFilter with *prob* set to 0.01 are congruent with each other, but show one difference that Gunnerales plus Dilleniales (BS = 0.75; 0.40; 0.50) rather than Gunnerales alone diverged first from the rest core eudicots. The trimAl ASTRAL tree with *st* set to 0.6 shows another different placement of Dilleniales, which is clustered within one clade (LPP = 0.56), in which, it is sister to Caryophyllales (LPP = 1.00) followed by asterids. The trimAl ASTRAL tree with *st* set to 0.7 shows another different placement, in which, Saxifragales is sister to Vitales (LPP = 0.44). Other ASTRAL trees show more variable topology. The alignmentFilter with *prob* set to 0.05 shows Gunnerales plus Dilleniales (LPP = 0.42), Berberidopsidales (LPP = 0.97) and Santalales (LPP = 0.67) sequentially diverged from the clade (LPP = 0.50) comprising the rest core eudicots. The rest core eudicots split into two clades. One comprises Caryophyllales and asterids (LPP = 0.76). In the other clade (LPP = 1.00), Saxifragales is sister to rosids (LPP = 0.85) followed by Vitales. The trimAl ASTRAL tree with *st* set to 0.8 shows a much more distinct topology, in which, Dilleniales alone diverged first from the rest core eudicots (LPP = 0.43), which in turn split into two clades. One clade (LPP = 0.34) includes Caryophyllales and asterids. In the other clade (LPP = 0.39), Gunnerales initially split from the rests (LPP = 0.77) which in turn diverged into rosids and a subclade. In the subclade (LPP = 0.49), Vitales (LPP = 0.40), Santalales (LPP = 0.95), Berberidopsidales and Saxifragales successively split from the rest(s). The relationships of other orders are more constant.


### Items

**Table S1** The RF distances between each pair of the plastid concatenation ML trees from the various alignment-filtering methods.

**Table S2** The RF distances between each pair of the nuclear concatenation ML trees from the various alignment-filtering methods.

**Table S3** The RF distances between each pair of the nuclear ASTRAL trees from the various alignment-filtering methods.

**Table S4** The RF distances between each pair of the simulation concatenation ML trees from the various alignment-filtering methods.

**Table S5** The RF distances between each pair of the simulation ASTRAL trees from the various alignment-filtering methods.

**Figure S1** The reduced trees based on the unfiltered plastid and nuclear data. (a0) plastid concatenation ML tree with support values from 100 bootstrap analyses. (b0) nuclear concatenation ML tree with support values from 100 bootstrap analyses. (c0) nuclear ASTRAL tree with LPP support values. These trees are used as comparisons for the respective set of trees (Figures S2–S4) based on the corresponding data filtered by the various alignment-filtering methods. The five Mesangiospermae clades except eudicots are highlighted in bold.

**Figure S2** The reduced plastid concatenation ML trees to show variable relationships of the 17 Mesangiospermae orders/superorders. The 11 trees are rooted with Austrobaileyales and nodal supports from 100 bootstraps are mapped. Trees a1–a11 are respectively from alignmentFilter (with parameter of *prob* set to 0.05, 0.01, 0.001 and 0.0001, respectively), Gblocks, HmmCleaner, TAPER and trimAl (with *st* set to 0.5, 0.6, 0.7 and 0.8, respectively). The five Mesangiospermae clades except eudicots are highlighted in bold.

**Figure S3** The reduced nuclear concatenation ML trees to show variable relationships of the 16 Mesangiospermae orders/superorders. The 12 trees are rooted with Austrobaileyales and nodal supports from 100 Bootstraps are mapped. Trees b1–b12 are respectively from alignmentFilter (with parameter of *prob* set to 0.05–0.0001, respectively), FasParser2, Gblocks, HmmCleaner, TAPER and trimAl (with *st* set to 0.5–0.8, respectively). The five Mesangiospermae clades except eudicots are highlighted in bold.

**Figure S4** The reduced nuclear ASTRAL trees to show variable relationships of the 16

Mesangiospermae orders/superorders. The 12 trees are rooted with Austrobaileyales and LPP support values are mapped. Trees c1–c12 are respectively from alignmentFilter (with parameter of *prob* set to 0.05–0.0001, respectively), FasParser2, Gblocks, HmmCleaner, TAPER and trimAl (with *st* set to 0.5–0.8, respectively). The five Mesangiospermae clades except eudicots are highlighted in bold.

**Figure S5** The practical differential performances of the various alignment-filtering methods on one of ambiguously-aligned regions in an empirical alignment. It is observed that alignmentFilter performs best while others have no (little) effect or delete whole columns along with well-aligned sites in a subset of sequences. The black cross symbols from the results of Gblocks and trimAl represent the contiguous sites as block that are deleted.

| | alignmentFilter -*prob*0.05 | alignmentFilter -*prob*0.01 | alignmentFilter -*prob*0.001 | alignmentFilter -*prob*0.0001 | Gblocks | HmmCleaner | TAPER | trimAl -*st*0.5 | trimAl -*st*0.6 | trimAl -*st*0.7 | trimAl -*st*0.8 |
|---|---|---|---|---|---|---|---|---|---|---|---|
| alignmentFilter-*prob*0.05 | 0 | | | | | | | | | | |
| alignmentFilter-*prob*0.01 | 6 | 0 | | | | | | | | | |
| alignmentFilter-*prob*0.001 | 6 | 4 | 0 | | | | | | | | |
| alignmentFilter-*prob*0.0001 | 4 | 2 | 2 | 0 | | | | | | | |
| Gblocks | 8 | 6 | 6 | 4 | 0 | | | | | | |
| HmmCleaner | 4 | 10 | 10 | 8 | 8 | 0 | | | | | |
| TAPER | 2 | 8 | 4 | 6 | 10 | 6 | 0 | | | | |
| trimAl-*st*0.5 | 4 | 8 | 8 | 6 | 6 | 4 | 6 | 0 | | | |
| trimAl-*st*0.6 | 8 | 12 | 8 | 10 | 6 | 8 | 6 | 4 | 0 | | |
| trimAl-*st*0.7 | 6 | 10 | 10 | 8 | 4 | 6 | 8 | 2 | 2 | 0 | |
| trimAl-*st*0.8 | 12 | 16 | 12 | 14 | 10 | 12 | 10 | 8 | 4 | 6 | 0 |

| | alignmentFilter -*prob*0.05 | alignmentFilter -*prob*0.01 | alignmentFilter -*prob*0.001 | alignmentFilter -*prob*0.0001 | FasParser2 | Gblocks | HmmCleaner | TAPER | trimAl -*st*0.5 | trimAl -*st*0.6 | trimAl -*st*0.7 | trimAl -*st*0.8 |
|---|---|---|---|---|---|---|---|---|---|---|---|---|
| alignmentFilter-*prob*0.05 | 0 | | | | | | | | | | | |
| alignmentFilter-*prob*0.01 | 2 | 0 | | | | | | | | | | |
| alignmentFilter-*prob*0.001 | 2 | 0 | 0 | | | | | | | | | |
| alignmentFilter-*prob*0.0001 | 10 | 10 | 10 | 0 | | | | | | | | |
| FasParser2 | 2 | 0 | 0 | 10 | 0 | | | | | | | |
| Gblocks | 2 | 4 | 4 | 12 | 4 | 0 | | | | | | |
| HmmCleaner | 0 | 2 | 2 | 10 | 2 | 2 | 0 | | | | | |
| TAPER | 0 | 2 | 2 | 10 | 2 | 2 | 0 | 0 | | | | |
| trimAl-*st*0.5 | 2 | 0 | 0 | 10 | 0 | 4 | 2 | 2 | 0 | | | |
| trimAl-*st*0.6 | 6 | 6 | 6 | 4 | 6 | 8 | 6 | 6 | 6 | 0 | | |
| trimAl-*st*0.7 | 16 | 16 | 16 | 6 | 16 | 18 | 16 | 16 | 16 | 10 | 0 | |
| trimAl-*st*0.8 | 24 | 24 | 24 | 14 | 24 | 26 | 24 | 24 | 24 | 18 | 16 | 0 |

| | alignmentFilter -*prob*0.05 | alignmentFilter -*prob*0.01 | alignmentFilter -*prob*0.001 | alignmentFilter -*prob*0.0001 | FasParser2 | Gblocks | HmmCleaner | TAPER | trimAl -*st*0.5 | trimAl -*st*0.6 | trimAl -*st*0.7 | trimAl -*st*0.8 |
|---|---|---|---|---|---|---|---|---|---|---|---|---|
| alignmentFilter-*prob*0.05 | 0 | | | | | | | | | | | |
| alignmentFilter-*prob*0.01 | 2 | 0 | | | | | | | | | | |
| alignmentFilter-*prob*0.001 | 2 | 4 | 0 | | | | | | | | | |
| alignmentFilter-*prob*0.0001 | 10 | 8 | 12 | 0 | | | | | | | | |
| FasParser2 | 4 | 6 | 2 | 14 | 0 | | | | | | | |
| Gblocks | 8 | 6 | 6 | 6 | 8 | 0 | | | | | | |
| HmmCleaner | 6 | 4 | 4 | 8 | 6 | 2 | 0 | | | | | |
| TAPER | 4 | 6 | 6 | 10 | 8 | 8 | 6 | 0 | | | | |
| trimAl-*st*0.5 | 4 | 2 | 6 | 6 | 8 | 4 | 2 | 4 | 0 | | | |
| trimAl-*st*0.6 | 10 | 8 | 12 | 12 | 14 | 10 | 12 | 12 | 10 | 0 | | |
| trimAl-*st*0.7 | 18 | 16 | 20 | 12 | 22 | 18 | 20 | 22 | 18 | 20 | 0 | |
| trimAl-*st*0.8 | 38 | 38 | 40 | 34 | 42 | 38 | 38 | 36 | 36 | 40 | 34 | 0 |

| | alignmentFilter-*prob*0.05 | alignmentFilter-*prob*0.01 | alignmentFilter-*prob*0.001 | alignmentFilter-*prob*0.0001 | FasParser2 | Gblocks | HmmCleaner | TAPER | trimAl-*st*0.5 | trimAl-*st*0.6 | trimAl-*st*0.7 | trimAl-*st*0.8 |
|---|---|---|---|---|---|---|---|---|---|---|---|---|
| alignmentFilter-*prob*0.05 | 0 | | | | | | | | | | | |
| alignmentFilter-*prob*0.01 | 0 | 0 | | | | | | | | | | |
| alignmentFilter-*prob*0.001 | 2 | 2 | 0 | | | | | | | | | |
| alignmentFilter-*prob*0.0001 | 2 | 2 | 0 | 0 | | | | | | | | |
| FasParser2 | 28 | 28 | 26 | 26 | 0 | | | | | | | |
| Gblocks | 28 | 28 | 26 | 26 | 0 | 0 | | | | | | |
| HmmCleaner | 18 | 18 | 16 | 16 | 20 | 20 | 0 | | | | | |
| TAPER | 8 | 8 | 6 | 6 | 22 | 22 | 14 | 0 | | | | |
| trimAl-*st*0.5 | 28 | 28 | 26 | 26 | 0 | 0 | 20 | 22 | 0 | | | |
| trimAl-*st*0.6 | 28 | 28 | 26 | 26 | 0 | 0 | 20 | 22 | 0 | 0 | | |
| trimAl-*st*0.7 | 24 | 24 | 22 | 22 | 4 | 4 | 16 | 18 | 4 | 4 | 0 | |
| trimAl-*st*0.8 | 10 | 10 | 8 | 8 | 22 | 22 | 14 | 6 | 22 | 22 | 18 | 0 |

| | alignmentFilter-*prob*0.05 | alignmentFilter-*prob*0.01 | alignmentFilter-*prob*0.001 | alignmentFilter-*prob*0.0001 | FasParser2 | Gblocks | HmmCleaner | TAPER | trimAl-*st*0.5 | trimAl-*st*0.6 | trimAl-*st*0.7 | trimAl-*st*0.8 |
|---|---|---|---|---|---|---|---|---|---|---|---|---|
| alignmentFilter-*prob*0.05 | 0 | | | | | | | | | | | |
| alignmentFilter-*prob*0.01 | 2 | 0 | | | | | | | | | | |
| alignmentFilter-*prob*0.001 | 4 | 2 | 0 | | | | | | | | | |
| alignmentFilter-0.0001 | 6 | 4 | 2 | 0 | | | | | | | | |
| FasParser2 | 4 | 2 | 4 | 6 | 0 | | | | | | | |
| Gblocks | 4 | 2 | 4 | 6 | 0 | 0 | | | | | | |
| HmmCleaner | 2 | 4 | 6 | 8 | 2 | 2 | 0 | | | | | |
| TAPER | 0 | 2 | 4 | 6 | 4 | 4 | 2 | 0 | | | | |
| trimAl-*st*0.5 | 4 | 2 | 4 | 6 | 0 | 0 | 2 | 4 | 0 | | | |
| trimAl-*st*0.6 | 4 | 2 | 4 | 6 | 4 | 4 | 6 | 4 | 4 | 0 | | |
| trimAl-*st*0.7 | 2 | 4 | 6 | 8 | 2 | 2 | 0 | 2 | 2 | 6 | 0 | |
| trimAl-*st*0.8 | 10 | 8 | 10 | 12 | 6 | 6 | 8 | 10 | 6 | 6 | 8 | 0 |

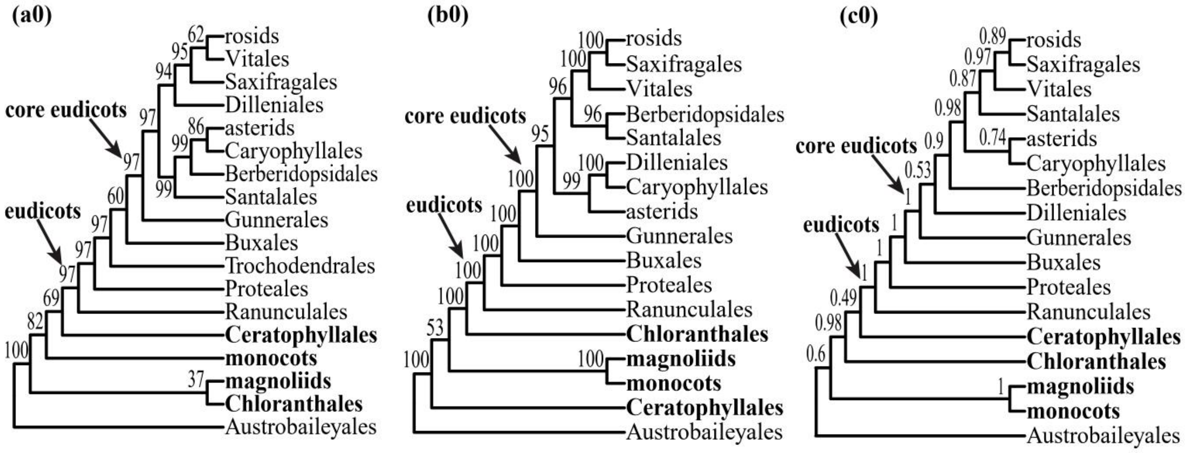
(a0)
core eudicots
eudicots
rosids
Vitales
Saxifragales
Dilleniales
asterids
Caryophyllales
Berberidopsidales
Santalales
Gunnerales
Buxales
Trochodendrales
Proteales
Ranunculales
Ceratophyllales
monocots
magnoliids
Chloranthales
Austrobaileyales
(b0)
core eudicots
eudicots
rosids
Saxifragales
Vitales
Berberidopsidales
Santalales
Dilleniales
Caryophyllales
asterids
Gunnerales
Buxales
Proteales
Ranunculales
Chloranthales
magnoliids
monocots
Ceratophyllales
Austrobaileyales
(c0)
core eudicots
eudicots
rosids
Saxifragales
Vitales
Santalales
asterids
Caryophyllales
Berberidopsidales
Dilleniales
Gunnerales
Buxales
Proteales
Ranunculales
Ceratophyllales
Chloranthales
magnoliids
monocots
Austrobaileyales

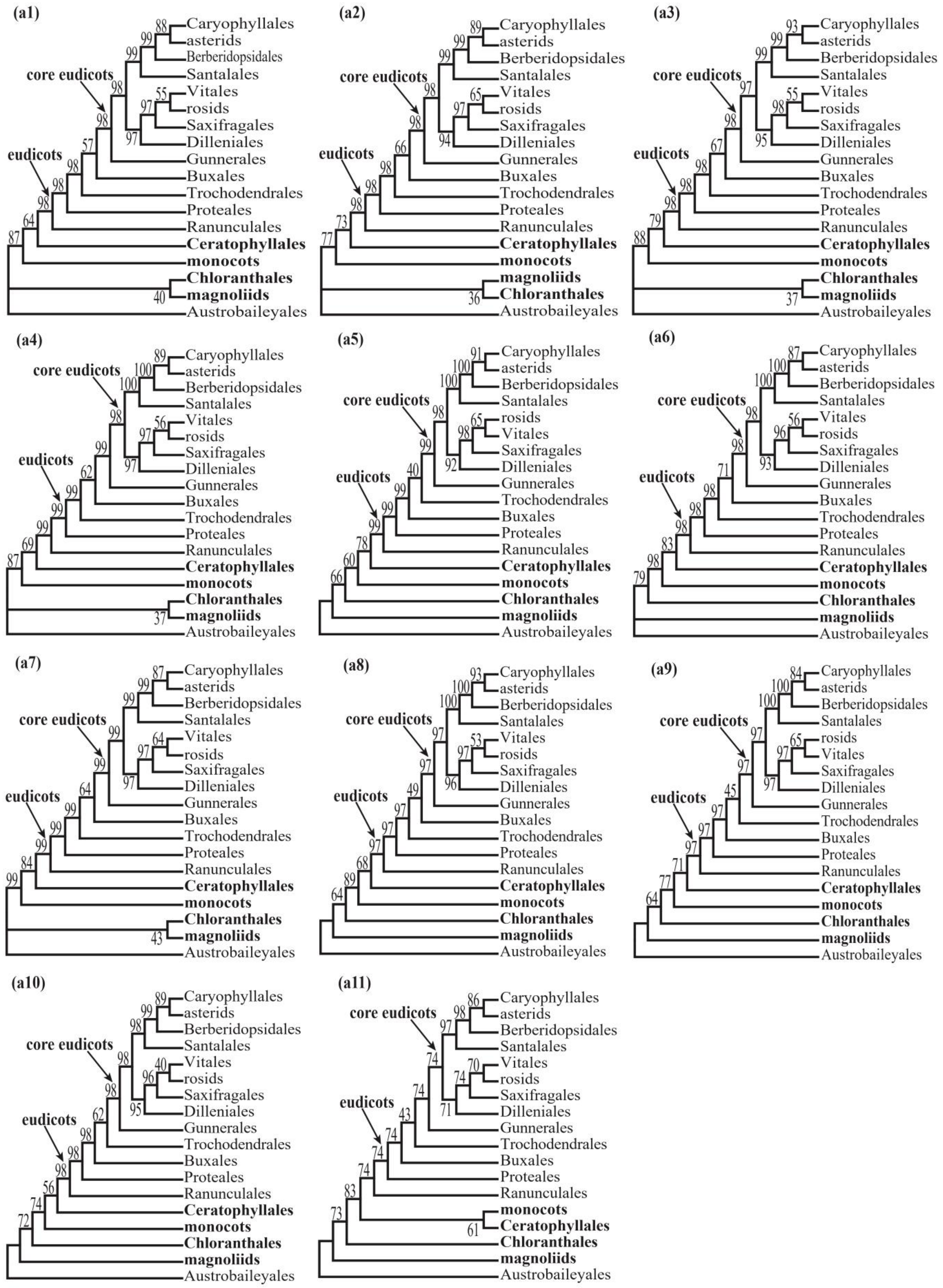
(a1)
core eudicots
eudicots
Caryophyllales
asterids
Berberidopsidales
Santalales
Vitales
rosids
Saxifragales
Dilleniales
Gunnerales
Buxales
Trochodendrales
Proteales
Ranunculales
Ceratophyllales
monocots
Chloranthales
magnoliids
Austrobaileyales
(a2)
core eudicots
eudicots
Caryophyllales
asterids
Berberidopsidales
Santalales
Vitales
rosids
Saxifragales
Dilleniales
Gunnerales
Buxales
Trochodendrales
Proteales
Ranunculales
Ceratophyllales
monocots
magnoliids
Chloranthales
Austrobaileyales
(a3)
core eudicots
eudicots
Caryophyllales
asterids
Berberidopsidales
Santalales
Vitales
rosids
Saxifragales
Dilleniales
Gunnerales
Buxales
Trochodendrales
Proteales
Ranunculales
Ceratophyllales
monocots
Chloranthales
magnoliids
Austrobaileyales
(a4)
core eudicots
eudicots
Caryophyllales
asterids
Berberidopsidales
Santalales
Vitales
rosids
Saxifragales
Dilleniales
Gunnerales
Buxales
Trochodendrales
Proteales
Ranunculales
Ceratophyllales
monocots
Chloranthales
magnoliids
Austrobaileyales
(a5)
core eudicots
eudicots
Caryophyllales
asterids
Berberidopsidales
Santalales
rosids
Vitales
Saxifragales
Dilleniales
Gunnerales
Trochodendrales
Buxales
Proteales
Ranunculales
Ceratophyllales
monocots
Chloranthales
magnoliids
Austrobaileyales
(a6)
core eudicots
eudicots
Caryophyllales
asterids
Berberidopsidales
Santalales
Vitales
rosids
Saxifragales
Dilleniales
Gunnerales
Buxales
Trochodendrales
Proteales
Ranunculales
Ceratophyllales
monocots
Chloranthales
magnoliids
Austrobaileyales
(a7)
core eudicots
eudicots
Caryophyllales
asterids
Berberidopsidales
Santalales
Vitales
rosids
Saxifragales
Dilleniales
Gunnerales
Buxales
Trochodendrales
Proteales
Ranunculales
Ceratophyllales
monocots
Chloranthales
magnoliids
Austrobaileyales
(a8)
core eudicots
eudicots
Caryophyllales
asterids
Berberidopsidales
Santalales
Vitales
rosids
Saxifragales
Dilleniales
Gunnerales
Buxales
Trochodendrales
Proteales
Ranunculales
Ceratophyllales
monocots
Chloranthales
magnoliids
Austrobaileyales
(a9)
core eudicots
eudicots
Caryophyllales
asterids
Berberidopsidales
Santalales
rosids
Vitales
Saxifragales
Dilleniales
Gunnerales
Trochodendrales
Buxales
Proteales
Ranunculales
Ceratophyllales
monocots
Chloranthales
magnoliids
Austrobaileyales
(a10)
core eudicots
eudicots
Caryophyllales
asterids
Berberidopsidales
Santalales
Vitales
rosids
Saxifragales
Dilleniales
Gunnerales
Trochodendrales
Buxales
Proteales
Ranunculales
Ceratophyllales
monocots
Chloranthales
magnoliids
Austrobaileyales
(a11)
core eudicots
eudicots
Caryophyllales
asterids
Berberidopsidales
Santalales
Vitales
rosids
Saxifragales
Dilleniales
Gunnerales
Trochodendrales
Buxales
Proteales
Ranunculales
monocots
Ceratophyllales
Chloranthales
magnoliids
Austrobaileyales

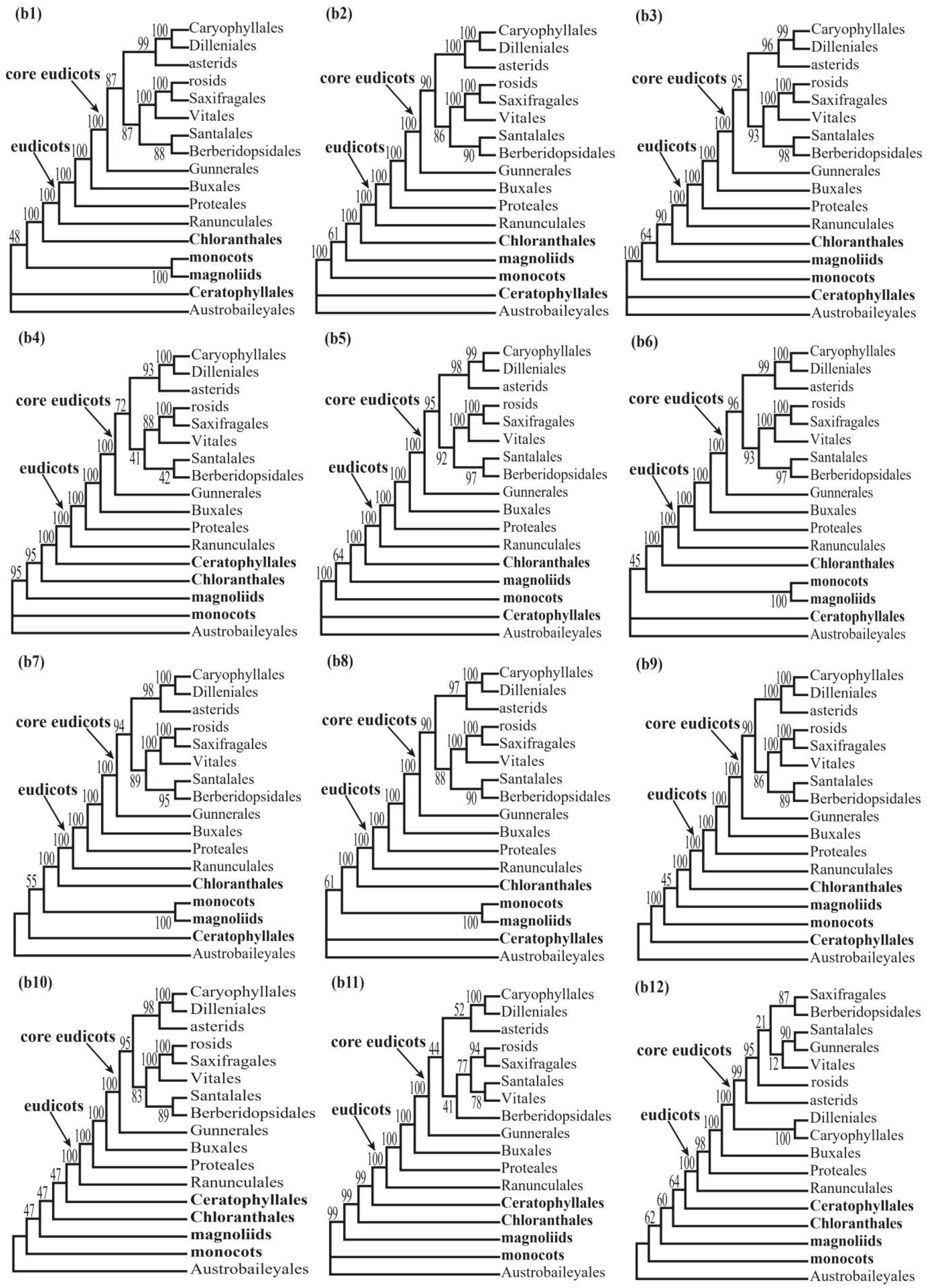
(b1)
core eudicots
eudicots
Caryophyllales
Dilleniales
asterids
rosids
Saxifragales
Vitales
Santalales
Berberidopsidales
Gunnerales
Buxales
Proteales
Ranunculales
Chloranthales
monocots
magnoliids
Ceratophyllales
Austrobaileyales
(b2)
core eudicots
eudicots
Caryophyllales
Dilleniales
asterids
rosids
Saxifragales
Vitales
Santalales
Berberidopsidales
Gunnerales
Buxales
Proteales
Ranunculales
Chloranthales
magnoliids
monocots
Ceratophyllales
Austrobaileyales
(b3)
core eudicots
eudicots
Caryophyllales
Dilleniales
asterids
rosids
Saxifragales
Vitales
Santalales
Berberidopsidales
Gunnerales
Buxales
Proteales
Ranunculales
Chloranthales
magnoliids
monocots
Ceratophyllales
Austrobaileyales
(b4)
core eudicots
eudicots
Caryophyllales
Dilleniales
asterids
rosids
Saxifragales
Vitales
Santalales
Berberidopsidales
Gunnerales
Buxales
Proteales
Ranunculales
Ceratophyllales
Chloranthales
magnoliids
monocots
Austrobaileyales
(b5)
core eudicots
eudicots
Caryophyllales
Dilleniales
asterids
rosids
Saxifragales
Vitales
Santalales
Berberidopsidales
Gunnerales
Buxales
Proteales
Ranunculales
Chloranthales
magnoliids
monocots
Ceratophyllales
Austrobaileyales
(b6)
core eudicots
eudicots
Caryophyllales
Dilleniales
asterids
rosids
Saxifragales
Vitales
Santalales
Berberidopsidales
Gunnerales
Buxales
Proteales
Ranunculales
Chloranthales
monocots
magnoliids
Ceratophyllales
Austrobaileyales
(b7)
core eudicots
eudicots
Caryophyllales
Dilleniales
asterids
rosids
Saxifragales
Vitales
Santalales
Berberidopsidales
Gunnerales
Buxales
Proteales
Ranunculales
Chloranthales
monocots
magnoliids
Ceratophyllales
Austrobaileyales
(b8)
core eudicots
eudicots
Caryophyllales
Dilleniales
asterids
rosids
Saxifragales
Vitales
Santalales
Berberidopsidales
Gunnerales
Buxales
Proteales
Ranunculales
Chloranthales
monocots
magnoliids
Ceratophyllales
Austrobaileyales
(b9)
core eudicots
eudicots
Caryophyllales
Dilleniales
asterids
rosids
Saxifragales
Vitales
Santalales
Berberidopsidales
Gunnerales
Buxales
Proteales
Ranunculales
Chloranthales
magnoliids
monocots
Ceratophyllales
Austrobaileyales
(b10)
core eudicots
eudicots
Caryophyllales
Dilleniales
asterids
rosids
Saxifragales
Vitales
Santalales
Berberidopsidales
Gunnerales
Buxales
Proteales
Ranunculales
Ceratophyllales
Chloranthales
magnoliids
monocots
Austrobaileyales
(b11)
core eudicots
eudicots
Caryophyllales
Dilleniales
asterids
rosids
Saxifragales
Santalales
Vitales
Berberidopsidales
Gunnerales
Buxales
Proteales
Ranunculales
Ceratophyllales
Chloranthales
magnoliids
monocots
Austrobaileyales
(b12)
core eudicots
eudicots
Saxifragales
Berberidopsidales
Santalales
Gunnerales
Vitales
rosids
asterids
Dilleniales
Caryophyllales
Buxales
Proteales
Ranunculales
Ceratophyllales
Chloranthales
magnoliids
monocots
Austrobaileyales

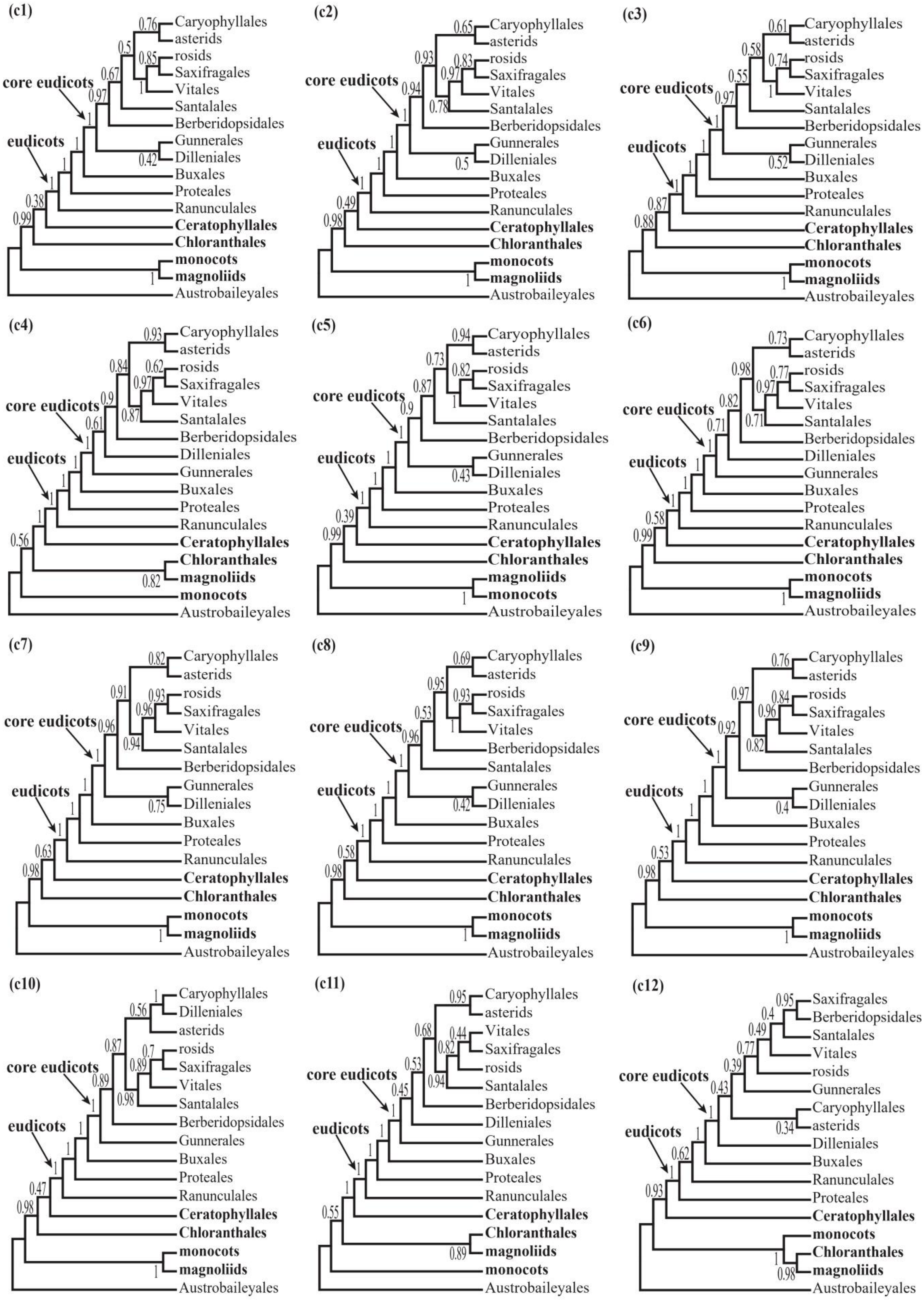
(c1)
core eudicots
eudicots
Caryophyllales
asterids
rosids
Saxifragales
Vitales
Santalales
Berberidopsidales
Gunnerales
Dilleniales
Buxales
Proteales
Ranunculales
Ceratophyllales
Chloranthales
monocots
magnoliids
Austrobaileyales
(c2)
(c3)
(c4)
(c5)
(c6)
(c7)
(c8)
(c9)
(c10)
(c11)
(c12)

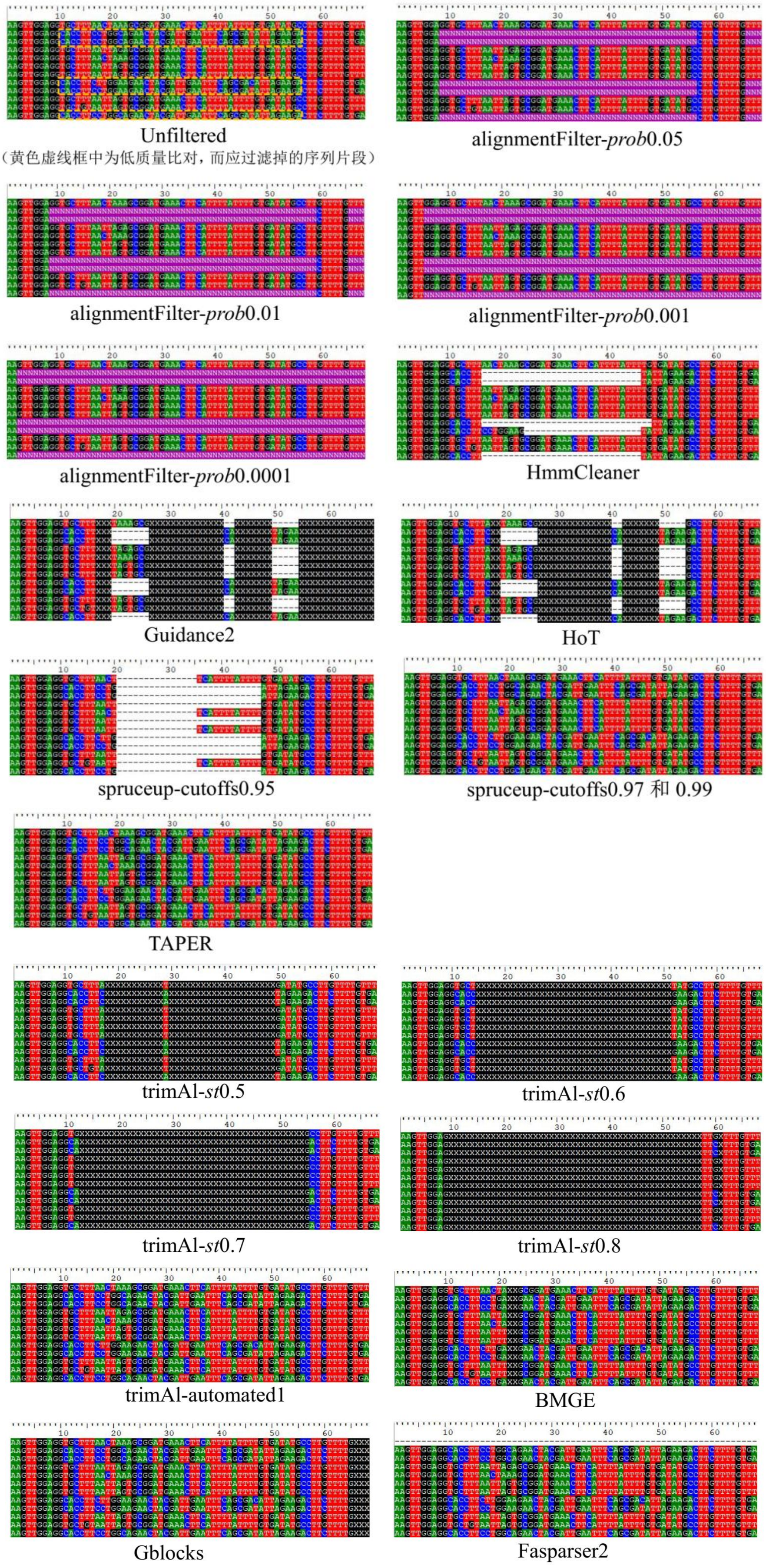
Unfiltered
（黄色虚线框中为低质量比对，而应过滤掉的序列片段）
alignmentFilter-*prob*0.05
alignmentFilter-*prob*0.01
alignmentFilter-*prob*0.001
alignmentFilter-*prob*0.0001
HmmCleaner
Guidance2
HoT
spruceup-cutoffs0.95
spruceup-cutoffs0.97 和 0.99
TAPER
trimAl-*st*0.5
trimAl-*st*0.6
trimAl-*st*0.7
trimAl-*st*0.8
trimAl-automated1
BMGE
Gblocks
Fasparser2
